# A Full Spectrum Fibroblast Profiling Panel Reveals Phenotypic and Metabolic Heterogeneity of Cancer-Associated Fibroblasts at the Single-Cell Level

**DOI:** 10.64898/2026.09.07.749987

**Authors:** Kevin Muñoz Forti, Marta Storl-Desmond, Sayana E. Isaac, Daniel Martínez, Michał Nizio, Shan Lin, Aniruddhsingh Solanki, Simon Schwörer

## Abstract

Cancer-associated fibroblasts (CAFs) exhibit extensive transcriptional and phenotypic heterogeneity and are highly plastic. Routine approaches to resolving CAF heterogeneity at the single-cell and protein levels are lacking, making it difficult to assess changes in CAF state in response to perturbations. Here, we present an optimized 28-marker spectral flow cytometry panel that uses unsupervised clustering to resolve CAF states across *in vivo* and *in vitro* models. Using this new fibroblast profiling panel (FPP), we show that LRRC15+ CAFs in murine pancreatic ductal adenocarcinoma are phenotypically distinct from canonical αSMA+ myofibroblasts, rather than representing a subset of this population. We find that marker expression is graded rather than discrete across clusters, indicating that CAF identity spans a continuum rather than fixed states. Metabolic profiling further shows that hypoxic CAFs were enriched for glucose uptake and an inflammatory phenotype. Using *in vitro* models, we further demonstrate how the FPP can resolve plasticity outcomes. In a 3D organoid co-culture model, direct contact between tumor cells and fibroblasts, but not organoid-derived factors alone, induces a stem-like CAF state also observed *in vivo*. In 2D, TGFβ and inflammatory/hypoxic stimuli each enrich distinct, pre-existing CAF states rather than generating new ones. These findings establish our FPP as a practical method for resolving fibroblast heterogeneity at the protein level in cancer and beyond.

## Introduction

Fibroblasts are mesenchymal cells present throughout normal tissue, where they maintain extracellular matrix (ECM) architecture and tissue homeostasis.^1,2^ Injury, inflammation, or other stress signals shift fibroblasts out of this basal, quiescent-like state into an activated phenotype characterized by increased contractility, heightened ECM deposition, and elevated production and secretion of growth factors and cytokines.^1^ This activation program typically supports wound healing and resolves once repair is complete; however, in tumors, it is sustained by signals from cancer cells, other stromal cell types, and the structural and metabolic changes within the tumor microenvironment (TME). This gives rise to cancer-associated fibroblasts (CAFs) that persist as a characteristic feature of solid tumors.^3^ CAFs are now recognized as major contributors to tumor progression, immune evasion, and treatment resistance, particularly in desmoplastic cancers such as pancreatic ductal adenocarcinoma (PDAC).^3–5^ Despite these pro-tumor functions, early efforts to deplete CAFs in PDAC in fact accelerated tumor progression.^6,7^ These seemingly contradictory findings have been explained, at least in part, by the substantial heterogeneity of CAFs.^8,9^

The concept of fibroblasts as heterogeneous cells was first described in 1977.^10^ Today, it is established that CAF heterogeneity is multifaceted, with cells displaying diverse cell-of-origin, tumor locations, gene expression profiles, and marker phenotypes.^8,9,11,12^ Diverse CAF transcriptional and phenotypic states may be attributed to distinct cell-of-origin, such as mesothelial cells,^13^ or niche-specific signals, such as TGFβ or hypoxia,^14,15^ reflecting the high plasticity of CAFs.^11,12^ Single-cell transcriptomic efforts have led to the emergence of a trichotomy model of CAF states that are conserved across tumor types: inflammatory CAFs (iCAFs) that express inflammatory cytokines, myofibroblastic CAFs (myCAFs) that express contractile genes and ECM modulators, and antigen-presenting CAFs (apCAFs) that express high levels of major histocompatibility complex (MHC) class II genes.^8,11^ Aside from transcriptomics, various single protein markers have been used to classify these CAFs phenotypically and functionally, and it was noted early on that a single marker alone will not identify all CAFs.^16^ αSMA has historically been used to define myofibroblasts in wound healing, fibrosis, and tumors.^17^ Later, it was used to describe activated pancreatic stellate cells,^18^ a CAF precursor population in PDAC, and was adopted as a general marker for CAFs,^19^ until it was specified as a myCAF marker for histological assessment of tumors.^20^ Notably, αSMA+ CAFs have tumor-restraining functions in PDAC.^6,21^ In contrast, FAPα+ CAFs are tumor-promoting in PDAC and other tumor types,^21–23^ in part by secreting immunosuppressive chemokines.^22^ FAPα has been associated with both the myCAF and iCAF state,^11^ and has been suggested to mark a heterogeneous population spanning the progenitor fibroblast-to-iCAF/myCAF continuum.^12^ LRRC15 marks a CAF population that promotes tumor growth across malignancies by suppressing anti-tumor immunity.^24,25^ LRRC15+ CAFs are induced by TGFβ signaling,^24,25^ suggesting a myCAF state; however, whether αSMA and LRRC15 denote the same CAF population remains unclear. Expression of CD105, a TGFβ co-receptor, distinguishes two CAF populations across tumor types, with CD105- CAFs restricting PDAC growth by supporting anti-tumor immunity.^26^ Interestingly, CD105- CAFs show high expression of MHC class II genes, suggesting they include apCAFs.^26^ In addition to these function-defining markers, other cell surface markers have been used to define CAF states by flow cytometry: Ly6C+/MHCII- CAFs were defined as iCAFs, and Ly6C-/MHCII+ CAFs were defined as apCAFs, with CAFs expressing neither of these markers denoted as myCAFs.^27^ This approach has since been widely adopted in the field. Recently, Ly6C-/MHCII- CAFs were further classified based on CD90 expression.^28^ Interestingly, Ly6C-/MHCII-/CD90- CAFs were sensitive to inhibition of pro-metastatic EGFR/ERBB2 signaling and had lower expression of both iCAF and myCAF transcriptional signatures compared to Ly6C-/MHCII-/CD90+ CAFs. These data indicate additional layers of heterogeneity beyond the established Ly6C/MHCII framework and highlight the need to include additional markers for CAF phenotyping. How different phenotypic markers relate to each other to define a CAF state remains to be investigated.

Mass cytometry using time-of-flight (CyTOF) offers greater dimensionality than conventional flow cytometry for phenotyping. It has been applied with a 42-marker panel to study mesenchymal heterogeneity in PDAC, resulting in the identification of CD105, as described above.^26^ Despite its dimensionality advantage, CyTOF’s destructive nature and high equipment costs have hindered its widespread adoption. While scRNA-sequencing is now widely used to profile heterogeneous CAF populations transcriptomically, it remains costly and time-consuming, limiting its use to profiling a limited number of precious samples in hypothesis-generating studies. There remains a need for affordable, practical, efficient, scalable, and reliable methods to assess the CAF state in routine hypothesis-testing experiments.

While advances in flow cytometry, such as the adoption of spectral flow cytometry and the development of novel fluorophores, have made panels of +40 colors possible, CAF biology has yet to benefit greatly from these technologies. This is due to various factors intrinsic to both CAFs and the technique. CAFs are relatively large cells compared to immune cells (2-10 times as big), challenging the most advanced optical systems, and they are found embedded in tissues rich in ECM, requiring extensive bespoke isolation protocols. Furthermore, there exists a paucity of fluorochrome-conjugated antibodies useful for studying CAF heterogeneity. This, coupled with the lack of formal consensus on which markers should be used to identify fibroblasts in general and CAFs in particular,^2^ has left this single-cell, non-destructive, quantitative, and high-throughput technology beyond the reach of CAF biology. Consequently, studies profiling CAF heterogeneity by flow cytometry have been limited to 10 markers or fewer,^23,27,29^ and the few markers in these panels to study CAFs directly were restricted to cell-surface markers. To bridge this gap and advance the study of CAF heterogeneity, we have designed, optimized, and validated a high-dimensional 28-marker spectral flow cytometry panel that leverages full-spectrum profiling to investigate fibroblast cell states and their dynamics in pancreatic cancer.

## Results

To evaluate the capabilities of spectral flow cytometry in combination with an established flow cytometry approach to resolve CAF states in PDAC,^27^ we established murine PDAC tumors via orthotopic injection of syngeneic KPC2-EYFP cells, followed by enzymatic dissociation using a method previously established to retain CAF surface marker expression.^24^ Samples were stained with CD326, CD31, CD45, PDPN, CD140α (PDGFRα), Ly6C, MHCII, CD90 and αSMA antibodies. For optimal spectral analysis and gating, we included autofluorescence (AF), single-antibody stained, and fluorescence-minus-one (FMO) controls (Fig. 1A). To detect αSMA, an intracellular protein, cells were fixed and permeabilized, a standard step in flow cytometry-based immune profiling but omitted from most CAF flow cytometry analyses. We used CD326, CD31, CD45, and EYFP to exclude epithelial cells, endothelial cells, immune cells, and cancer cells, respectively, followed by positive selection for PDPN and/or CD140α to define CAFs, and Ly6C/MHCII analysis to define CAF subsets (Fig. 1B).^27^ Consistent with prior work,^27^ Ly6C- /MHCII- CAFs were the most prominent CAF subset identified in this model (Fig. 1C). As this subset of CAFs is presumed to represent myCAFs,^27^ we further analyzed αSMA and CD90 expression in Ly6C-/MHCII- CAFs to test whether these cells are *bona fide* myofibroblasts, according to historical and more recent definitions.^17,28^ This analysis resolved four populations, including a prominent subset of cells expressing neither αSMA nor CD90 (Fig. 1C), suggesting that they may not represent myofibroblasts. These data indicate that the most widely used flow cytometry strategy, even when complemented with two additional markers, incompletely resolves CAF heterogeneity in PDAC.

**Figure 1.**
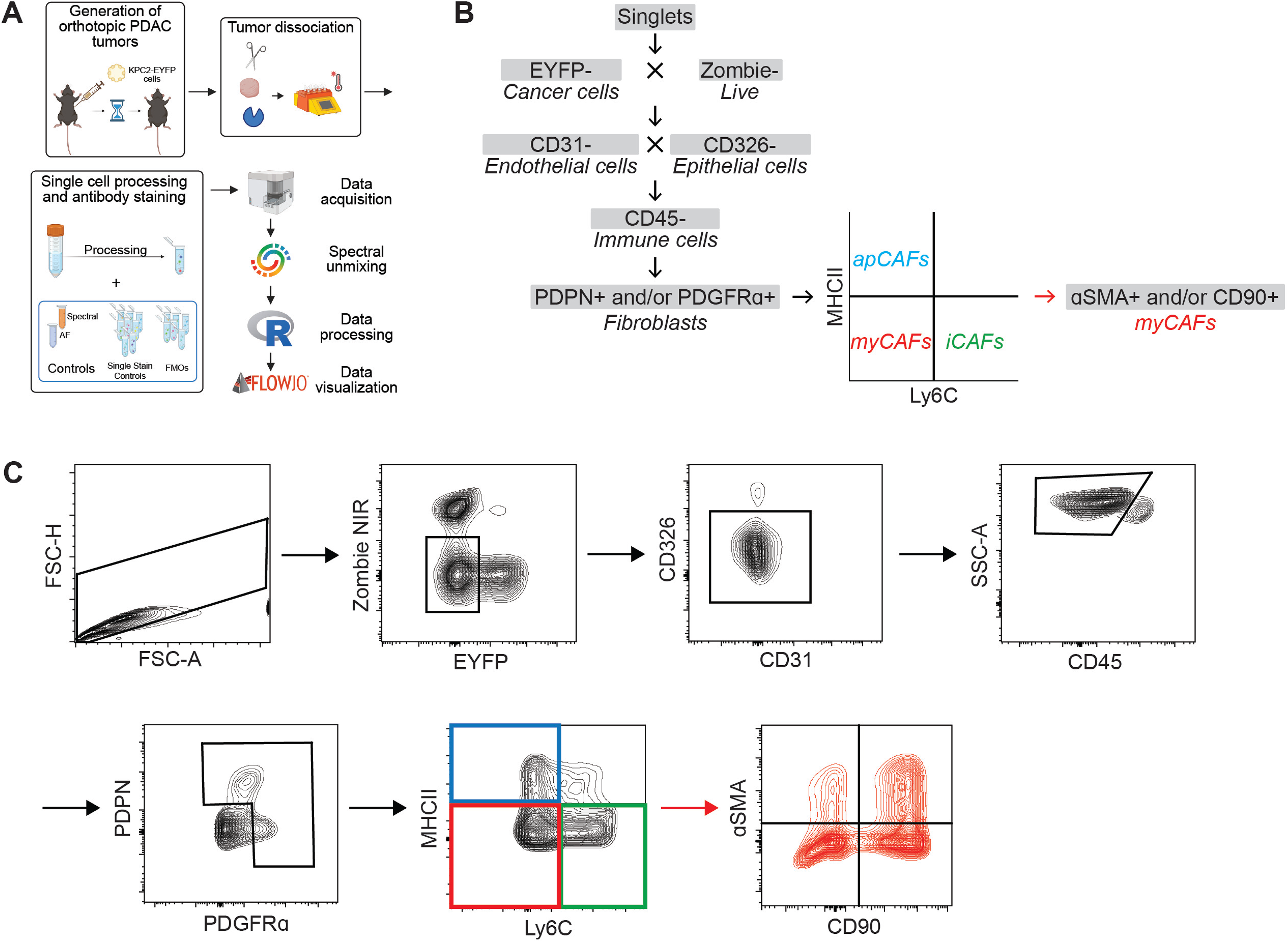
Spectral flow cytometry-based identification of previously reported cancer- associated fibroblasts (CAFs) states. **(A)** Overview of workflow. Single-cell suspensions were prepared from PDAC tumor tissues using enzymatic digestion and physical dissociation. Cells were stained for initial flow cytometry analysis to identify CAFs based on phenotypic markers. **(B, C)** Schematic (B) and contour plots (C) representations of the gating strategy to identify CAF subpopulations. CAFs were identified as negative/low for Zombie NIR, negative for CD45, CD326, CD31, and EYFP expression, and positive for podoplanin (PDPN) and/or platelet- derived growth factor receptor alpha (PDGFRα). CAFs were further subset into MHC II+Ly6C- antigen-presenting CAFs (apCAFs), MHC II-Ly6CLo/- myofibroblastic CAFs (myCAFs), and MHC II-Ly6C+ inflammatory CAFs (iCAFs). αSMA and/or CD90 were used to further define the myCAF subpopulation.

To overcome these limitations, we designed a new full-spectrum profiling panel, which we termed fibroblast profiling panel (FPP). To ease adoption of the FPP by laboratories, we based it on the widely used markers described above. We included additional pan-fibroblast markers, effector molecules, and markers of fibroblast phenotypic and functional heterogeneity across tumor types, focusing on commercially available antibody-fluorophore conjugates and off-the-shelf reagents. This included markers such as CD140β/PDGFRβ, CD29/ITGB1, LRRC15, FAPα, S100A4/FSP1, CD274/PD-L1, CD73/NT5E, CD74, CD34, Ly6A/Sca1, IL6, and CXCL1 (Table 1). This core, 26-marker FPP has a Complexity Index of 16.13 (Sup. Fig. 1A), which is comparable to validated commercial full-spectrum panels of similar scale (e.g., Cytek’s 24-color mouse immunophenotyping panel, Complexity Index=11.7) and within the range reported for other validated spectral panels (12.82-31.16),^30^ supporting adequate spectral resolution. Each antibody-fluorophore conjugate was titrated to determine optimal concentration (Sup. Fig. 1B, C), and its spectral signature compared between cells from murine PDAC tumors and compensation beads for optimal spectral unmixing (Sup. Fig. 2). To test whether the additional markers included in the FPP improve resolution of distinct fibroblast subpopulations within Zombie-/CD45-/CD326-/CD31-/EYFP-, PDPN+ and/or CD140α+ CAFs beyond the established pan-CAF and CAF state markers PDPN, CD140α, Ly6C and MHCII, we iteratively tested the homogeneity of variances of cell clusters as a function of the number of markers using Bartlett’s test (Sup. Fig. 3). Each marker combination increased Bartlett’s chi-square statistic (Sup. Fig. 3A), indicating increased cluster resolvability. Notably, Bartlett’s chi-square statistic showed a shallower slope beyond a combination of 14 CAF markers, with CD34 and Ly6A as the last markers incorporated, indicating continued but smaller contribution of these markers to cluster resolvability (Sup. Fig. 3A, B). Conversely, adding CXCL-1, αSMA, CD140β, CD29, FSP1, CD73, and CD90 to the established four markers had the largest impact on cluster resolvability (Sup. Fig. 3B), suggesting these markers capture maximal non-redundant covariance within the phenotypic space. Together, these data support the inclusion of the selected additional protein markers in the FPP.

**Table 1.**
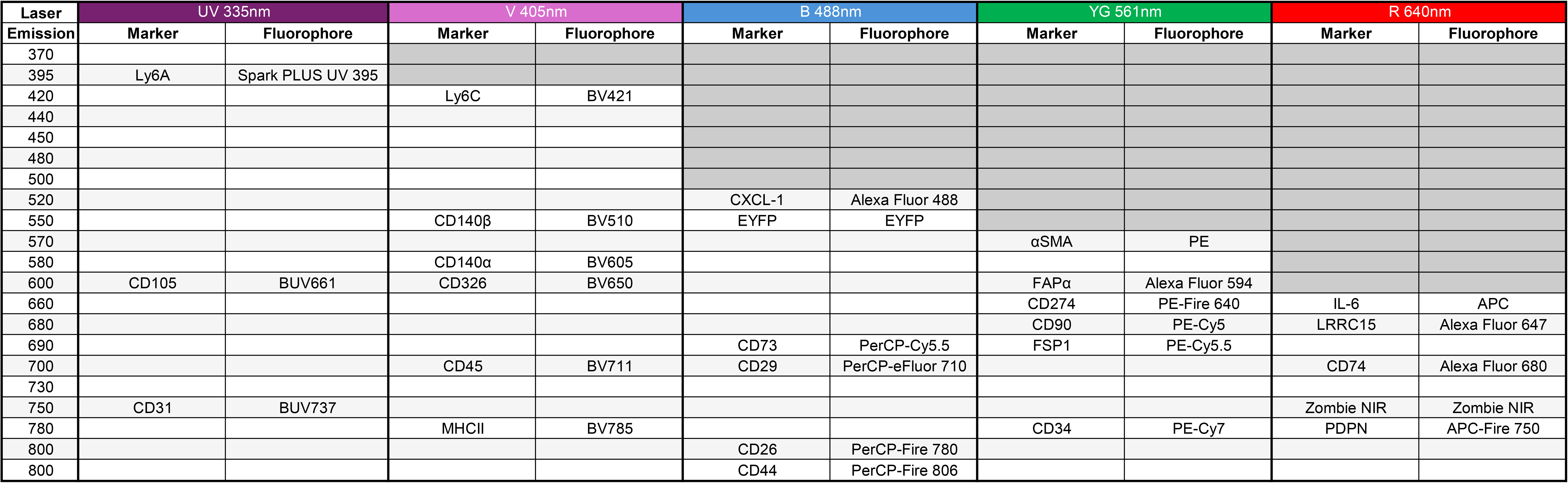
Fluorophore-marker pairings for core fibroblast state profiling panel. 26-marker panel excluding AF signatures and metabolic probes 2NBDG and Pimo.

Beyond protein markers, heterogeneous populations of CAFs have been described based on metabolic properties, including localization in hypoxic regions^15,31^ and glucose uptake.^32,33^ To account for metabolic heterogeneity of CAFs, we also included two metabolic probes – the hypoxia marker pimonidazole (Pimo) and the fluorescent glucose analog 2-NBDG – in our FPP, resulting in a total of 28 markers. To demonstrate the utility of the FPP, we generated KPC organoid-derived orthotopic PDAC tumors that closely recapitulate key features of human PDAC and exhibit hypoxic niches.^15,34^ Before the experimental endpoint, Pimo was administered *in vivo*, followed by single-cell dissociation and *ex vivo* treatment with 2-NBDG (Fig. 2A). Following data acquisition, unmixing and processing, we gated for CAFs using negative and positive selection described above, and visualized the high dimensional flow cytometry data as Uniform Manifold Approximation and Projection (UMAP) (Fig. 2A). The 17 CAF-specific markers and two metabolic markers of the FPP were projected on a UMAP from a representative tumor (Fig. 2B). To characterize CAF heterogeneity, we used PhenoGraph for clustering, as it makes no prior assumption about the number or size of subpopulations. When applied across three independent tumors, PhenoGraph identified 16-19 clusters per tumor, which were visualized using UMAP (Fig. 2C; Sup. Fig. 4A, B) and annotated by unsupervised hierarchical clustering of marker expression (Fig. 2D; Sup. Fig. 4C, D). Most markers displayed graded expression in several CAF clusters, indicating a spectrum of phenotypic states (Fig. 2B-D; Sup. Fig. 4A-D).

**Figure 2.**
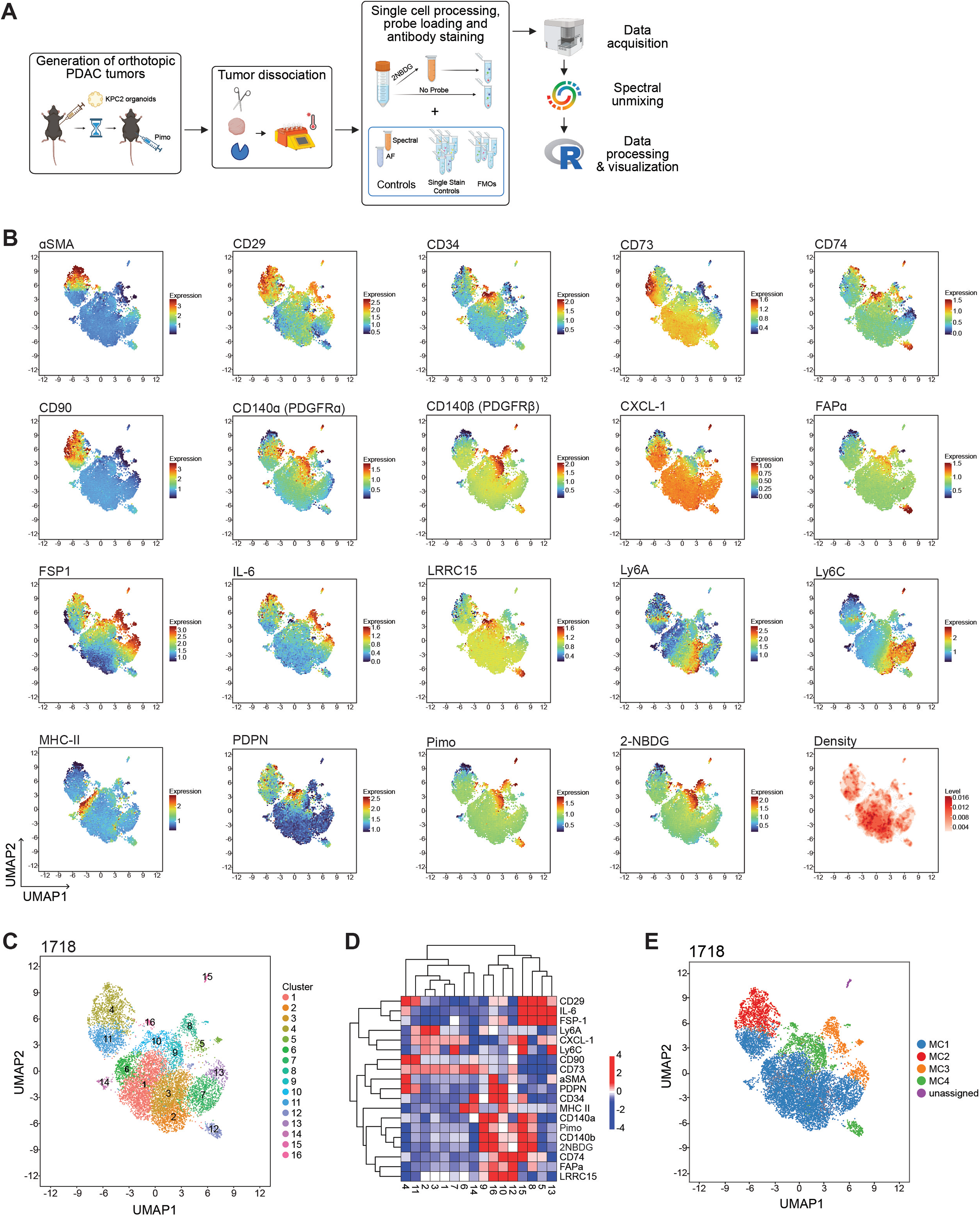
Implementation of 28-marker CAF panel in murine PDAC. **(A)** Overview of workflow. Tumor-bearing mice were injected with Pimo 1 hour before euthanasia. Single-cell suspensions were prepared from PDAC tumor tissues using enzymatic digestion and physical dissociation. Cells were incubated with 2-NBDG *ex vivo* and then stained for flow cytometry analysis. **(B)** Representative UMAPs of events defined as CAFs using the core fibroblast state profiling panel, with expression of inclusion markers and metabolic probes Pimo and 2-NBDG projected. Tumor 1718 is shown. **(C)** Representative UMAP with PhenoGraph clustering projected. Tumor 1718 is shown. **(D)** Heatmap of marker expression across PhenoGraph clusters with hierarchical clustering. Tumor 1718 is shown. **(E)** Representative UMAP with metaclustering based on Euclidean distance. Tumor 1718 is shown.

To identify recurring CAF states across replicate tumors, we used the following strategies: 1) metaclustering (MC) based on the Euclidean distance from the hierarchical clustering of PhenoGraph clusters, allowing us to group clusters similar in marker expression within each tumor; 2) pairwise correlation of individual PhenoGraph cluster combinations to identify a tumor cluster’s best correlated clusters across the other two tumors; 3) correlation of all unique marker pairs across tumors. This analysis revealed four MCs within each tumor (Fig. 2E; Sup. Fig. 4E, F), with only one small (<1% of cells) PhenoGraph cluster per tumor that did not match the marker expression profiles of the other tumors and therefore was not assigned to an MC. MC2 displayed high expression of CD90, CD29, and αSMA, but low levels of LRRC15, FAPα, MHCII, and 2-NBDG, indicative of a classical, contractile myCAF state.^20,21^ Within MC2, clusters c4/c10/c4 (tumors 1719/LET-888/1736) showed the highest correlation across tumors (*r*=0.827). In addition, αSMA and CD90 expression highly correlated across tumors (Sup. Fig. 5A, B), which was also observed in an independent scRNA-sequencing dataset from the KPC autochthonous tumor model^27^ (Sup. Fig. 5C). MC3 was characterized by high levels of IL6, FSP1, Pimo, and 2-NBDG, but low LRRC15, CD90, and CD73 expression. This marker profile aligns with the iCAF state enriched in hypoxic niches and characterized by high IL6 expression.^15,20^ Within MC3, clusters c8/c15/c1 were most similar across tumors (*r*=0.931). Pimo and 2-NBDG correlated across tumors (Sup. Fig. 5B), consistent with the known relationship between hypoxia and glucose uptake.35 In addition, an anticorrelation between IL6 and LRRC15 was observed (Sup. Fig. 5B), which was confirmed in KPC tumors at the transcriptional level (Sup. Fig. 5C). Similarly, we found a correlation between Pimo levels and both CD140α and Ly6C (Sup. Fig. 5B), which are expressed at higher levels in iCAFs than myCAFs.^27^ At the mRNA level, expression of these markers correlated with a hypoxic gene expression signature in CAFs from KPC mice (Sup. Fig. 5C). MC4 showed high expression of LRRC15, FAPα and CD74 but only moderate expression of αSMA and CD90, altogether indicative of an alternative, LRRC15+/FAPα+ myCAF state.^24,25^ LRRC15 and FAPα were highly correlated in orthotopic tumors (Sup. Fig. 5B), consistent with autochthonous PDAC tumors (Sup. Fig. 5C). Notably, the highest correlated clusters c12/c14/c17 (*r*=0.757) within MC4 across tumors were anticorrelated in their marker expression profile with the contractile myCAF clusters c4/c10/c4 (*r*=-0.64). Interestingly, this cluster combination (c12/c14/c17) occupied a region of UMAP space separate from the remaining PhenoGraph clusters assigned to MC4 (Fig. 2E; Sup. Fig. 4E, F). Finally, MC1, the largest metacluster identified, comprised several closely related PhenoGraph clusters based on Euclidean distance that showed no reliably elevated markers across tumors, indicating an overall less polarized state than MC2, MC3, and MC4. Within MC1, two related states were consistently found: a Ly6A-high, Ly6C-moderate state, with clusters c3/c5/c2 (*r*=0.819) as the most correlated across tumors, and a Ly6C-high, FSP1-moderate, Ly6A-low state represented by clusters c7/c8/c18 (*r*=0.638). Ly6A and Ly6C protein and mRNA expression were highly correlated in orthotopic and autochthonous PDAC models, respectively (Sup. Fig. 5B, C), and this marker combination has been used to denote Pi16+ universal fibroblasts.^36^

The above data indicate that our FPP can resolve several highly polarized and less polarized CAF states in murine PDAC tumors. CAF states have been modeled *in vitro* using 3D cultures of pancreatic stellate cells (PSCs) alone or together with PDAC organoids or their secreted factors via conditioned media (CM).^20^ To test if the FPP is similarly capable of resolving CAF states in this model, we applied it to cultures of two independent PSC lines (PSC1d, PSC-HB) in 2D, 3D, 3D KPC organoid co-cultures, or 3D with KPC organoid CM (Fig. 3A). PSC1d have been immortalized by spontaneous outgrowth and described,^15^ and newly generated PSC-HB were SV40 immortalized and their fibroblast nature validated (Sup. Fig. 6A). Given that the culture conditions use here don’t feature hypoxia, metabolic probes were omitted from the FPP, but recently described, additional protein markers for CAF state included, such as CD26/DPP4 and CD105/ENG.^23,26^ PSCs were identified as PDPN+/CD326- in co-cultures (Fig. 3B). UMAPs segregated PSCs between 2D/3D and 3D co-culture/CM (Fig. 3C; Sup. Fig. 6B), indicating that our FPP can resolve state changes induced by the presence of organoids or organoid-derived factors. PhenoGraph identified 13 and 15 clusters, respectively, that display unique marker expression profiles (Fig. 3D, E; Sup. Fig. 6C, D), mapped to different culture conditions, and were highly reproducible across replicates (Fig. 3F; Sup. Fig. 6E). Metacluster analysis identified five shared MCs between PSC lines plus two additional, line-specific MCs (Fig. 3G, H; Sup. Fig. 6F, G). Culture conditions drove dominance of specific MCs (ANOVA interaction *p*<0.0001) (Fig. 3H; Sup. Fig. 6G). While cell PSC line-specific differences were observed in MC1, MC4 and MC5 were composed exclusively of cells from the 2D and 3D groups (Fig. 3H; Sup. Fig. 6G). Both MC4 and MC5 cells showed high expression of CD26, LRRC15, and moderate expression of αSMA, indicative of a myofibroblastic nature. MC4 displayed low expression of CD105 and CD74, while MC5, enriched in 2D (*p*<0.05), had high expression of FAPα, CD90, and CD105 and was negative for MHCII. These data are consistent with an established model in which the supraphysiological stiffness of 2D culture drives PSCs into a myofibroblastic state, a shift that is dampened in 3D culture.^20^ MC3 was composed exclusively of cells from the 3D co-culture and 3D CM groups, with a higher proportion from the CM group than from the co-culture setting (*p*<0.05). Compared with other MCs, MC3 cells expressed elevated levels of CD140β, IL6, CD73, and MHCII, and low levels of CD26, collectively indicating a more inflammatory state; however, αSMA and FAPα expression remained moderate. Analysis of PhenoGraph clusters within MC3 suggested a continuum from a purer inflammatory state (c7 in PSC1d, c2/13 in PSC-HB) to co-expression of myofibroblast markers (c1/c2 in PSC1d, c3/c12 in PSC-HB), consistent with organoid-derived factors capable of inducing both inflammatory and myofibroblast features in PSCs.^14,28^ Finally, MC2 was composed exclusively of cells from the co-culture treatment group. It comprised one PhenoGraph cluster per line (c10 in PSC1d, c5 in PSC-HB), although PSC1d’s c5, part of the line-specific MC7, showed the highest similarity to PSC-HB’s c5 (r=0.8166) at the individual-cluster level. These clusters were characterized by high expression of Ly6A and Ly6C and moderate CD105 expression, with all other markers low, reminiscent of the Ly6A/Ly6C-high clusters c3/c5/c2 in PDAC tumors (Fig. 2D; Sup. Fig. 4C, D).

**Figure 3.**
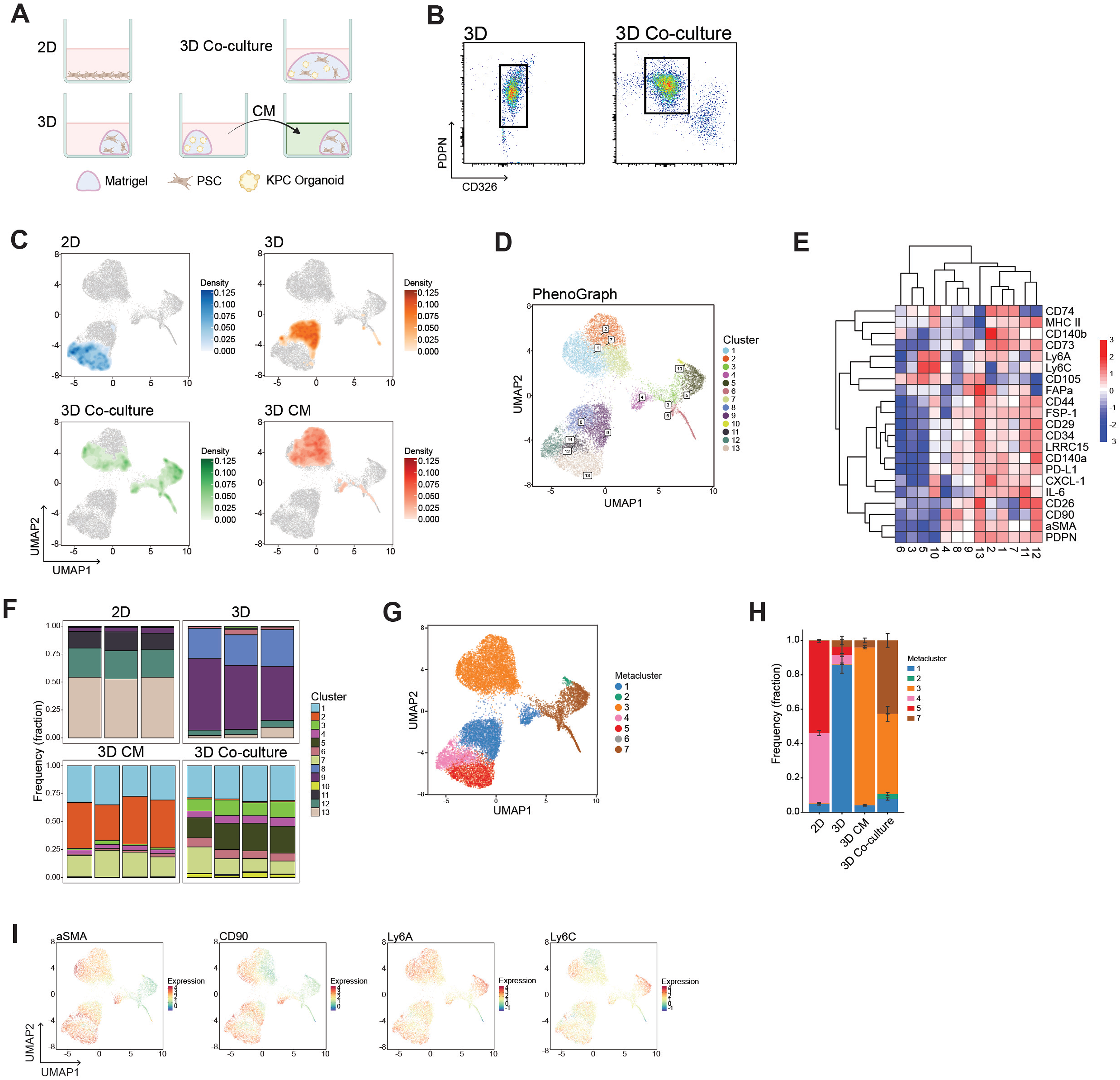
PDAC organoid co-culture exhibits differential direct and indirect effects on PSC1d fibroblast cell state. **(A)** Workflow schematic of 3D culture model. **(B)** Representative gating demonstrating effective separation of PSCs (PDPN+/CD326-) from organoids (PDPN-/CD326+). **(C)** UMAPs colored by cell density, faceted by culture condition. **(D)** UMAP with PhenoGraph clustering projected. **(E)** Heatmap of marker expression across PhenoGraph clusters with hierarchical clustering. **(F)** Bar charts of composition of each PhenoGraph cluster as a function of sample grouping. **(G)** UMAP with metaclustering based on Euclidean distance. **(H)** Bar charts of composition of each metacluster as a function of sample grouping. *n*=3 (2D/3D), *n*=4 (3D CM, 3D Co-culture) biological replicates. Data show mean +/- SD. **(I)** UMAPs with the expression of select CAF markers projected.

The above data demonstrate FPP’s capacity to resolve fibroblast states in physiologically relevant 3D culture models. While 2D culture models suffer from artificially high stiffness, which drives myofibroblastic properties in fibroblasts, they remain widely used in the field, and their relevance can be improved by incorporating tumor-relevant environmental factors, such as hypoxia.^15^ To demonstrate the utility of the FPP in this modality, we treated PSCs with TGFβ to further promote the myofibroblastic program, or with a combination of IL1α/TNFα/LIF (hereafter referred to as “cytokines”) and culture in 1% O2 (“hypoxia”), to promote an inflammatory program^15^ (Fig. 4A). Cells were resolved by culture condition on the UMAP projection, particularly TGFβ and Cytokines+Hypoxia conditions clustered separately (Fig. 4B). PhenoGraph identified 19 clusters with unique marker profiles (Fig. 4C, D) that were differentially but reproducibly enriched across culture conditions (Fig. 4E). Metacluster analysis identified six MCs (Fig. 4F), with MC3 accounting for <2.5% of cells within the different culture conditions (Fig. 4G) and was therefore not considered for further analysis. Notably, the remaining MCs were split into two broad programs based on expression of Ly6C, Ly6A, CD34, and CD140α, markers of both universal and inflammatory fibroblast states.^27,36^ These markers were low in MC1 and moderate to high in MC2 (Fig. 4D, H), indicating differences in stem-like and/or inflammatory properties. These programs were further split into two (1a, 1b) or three (2a, 2b, 2c) MCs driven by LRRC15 and MHCII expression (Fig. 4D, H). As with the 3D model (Fig. 3), culture conditions shaped the dominance of a particular MC (ANOVA interaction, *p*<0.0001) of the PSCs from the 2D model (Fig. 4G). While all MCs were present at baseline (vehicle), reflecting PSC heterogeneity, TGFβ treatment collapsed MCs 2a/2b/2c (*p*<0.0001) and inflated MCs 1a/1b (*p*<0.001), indicating a highly polarized state (Fig. 4G). The largest single cluster within the TGFβ condition, c10 (MC1a), showed a strong myofibroblastic profile (αSMA-high, CD90-high, LRRC15-high, CD26-high, and MHCII-negative). Interestingly, TGFβ also produced two large LRRC15-low and MHCII-moderate clusters within MC1b (c8, c14) harboring high IL6 and moderate FAPα expression. In contrast to TGFβ, Cytokine+Hypoxia treatment contributed fewer cells to the MC 1a/1b states (*p*<0.001) but increased cells within the MC2c state (*p*<0.001) compared to vehicle, overall increasing the proportion of cells with a universal/inflammatory profile (Fig. 4G). Specifically, highly abundant MC2c cells were LRRC15- low, MHCII-moderate, and IL6- and CXCL1-moderate to high, consistent with an inflammatory state. Taken together, these data demonstrate FPP’s capacity to resolve PSC heterogeneity at baseline and in response to environmental stimuli, showing how environmental factors drive highly heterogeneous PSCs into fewer but highly polarized states.

**Figure 4.**
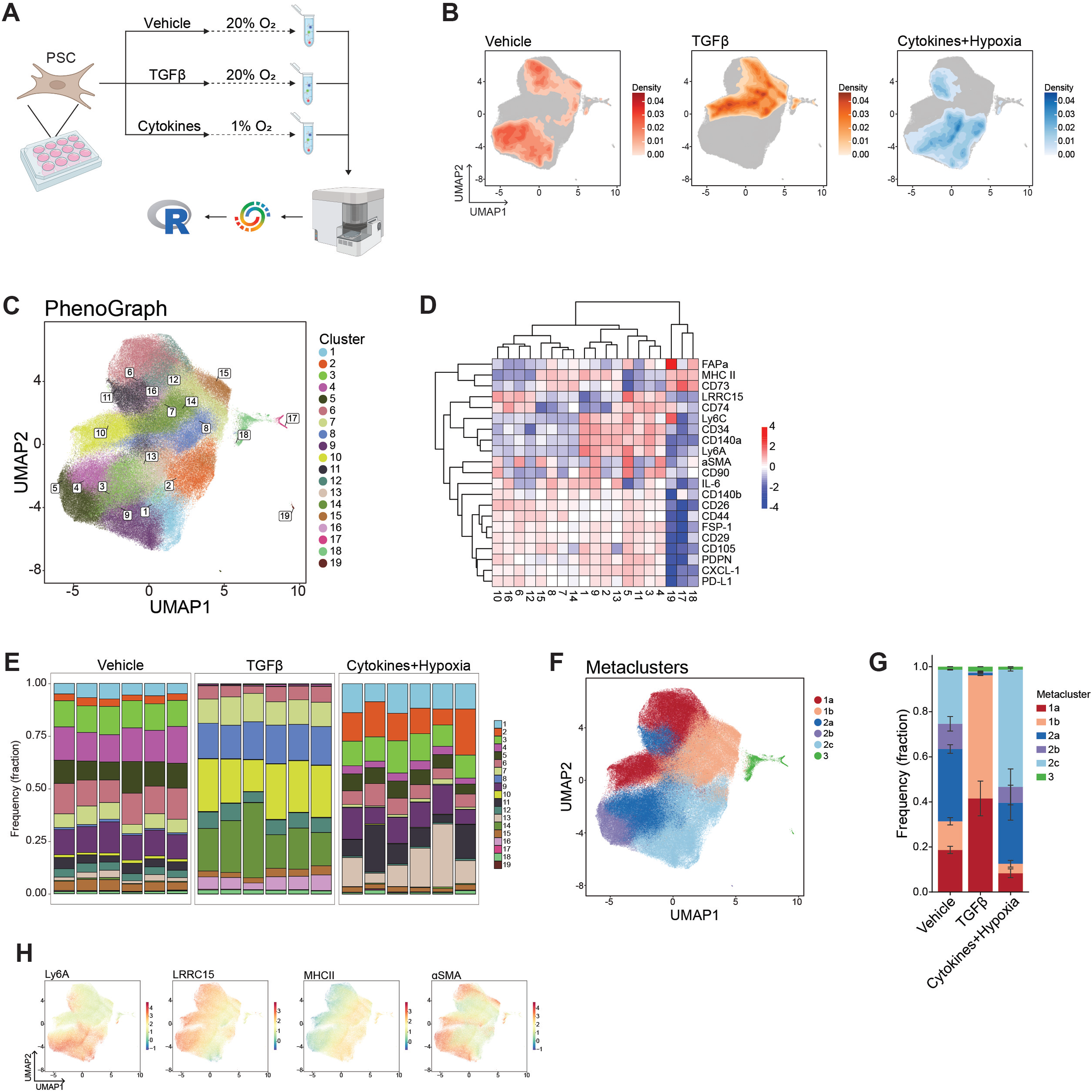
Fibroblast state profiling recapitulates the effects of fibroblast activating factors. **(A)** Workflow schematic. PSCs were cultured under conditions that promote the iCAF state (cocktail of cytokines and culture in hypoxia) and the myCAF state (treatment with TGFβ). **(B)** UMAPs colored by cell density, faceted by treatment group. **(C)** UMAP with PhenoGraph clustering projected. **(D)** Heatmap of marker expression across PhenoGraph clusters with hierarchical clustering. **(E)** Bar charts of composition of each PhenoGraph cluster as a function of sample treatment. **(F)** UMAP with metaclustering based on Euclidean distance. **(G)** Bar charts of composition of each metacluster as a function of sample grouping. *n*=6 biological replicates. Data represent mean +/- SD. **(H)** UMAPs with the expression of select CAF markers projected.

## Discussion

CAFs are key orchestrators of the TME, regulating ECM architecture, immune responses, and tumor growth.^2^ Their extensive heterogeneity is well established at the transcriptional level; however, tools to adequately resolve this heterogeneity at the protein level in single cells have lagged. Conventional flow cytometry strategies leave a substantial fraction of CAFs uncharacterized (Fig. 1). Here, we present an optimized, high-dimensional spectral flow cytometry pipeline that integrates unsupervised clustering and correlation networks to systematically resolve CAF states across various *in vivo* and *in vitro* experimental models.

In contrast to single-cell transcriptomics and mass cytometry, the FPP is a flexible, efficient, and affordable tool for studying fibroblast and CAF heterogeneity in routine laboratory experiments while maintaining single-cell resolution. We designed our FPP based on a skeleton panel resembling routinely used CAF markers (Fig. 1)^27^ and added commercially available, pre-conjugated antibodies. To isolate true phenotypic signatures from technical noise, we implemented a rigorous quality control workflow. Doublet gating and automated anomaly detection via FlowAI^37^ minimized instrument artifacts. Tissue autofluorescence was extracted as an independent spectral signature during unmixing, using unstained cells from each tumor and cell type, preventing misassignment of autofluorescent signal to dim markers. Equalizing sample representation through downsampling ensured that cohort-level analyses were not biased by individual sample cell counts, establishing a clean biological starting point. To address the practical challenges of designing streamlined yet information-preserving cytometry panels for CAF immunophenotyping, we implemented a combinatorial marker optimization framework based on Bartlett’s test of sphericity (Sup. Fig. 3). A larger test statistic reflects greater shared covariance structure, the hallmark of a biologically informative, non-redundant marker panel. This approach provides a statistically principled, data-driven alternative to subjective panel reduction strategies based solely on marker expression frequency or researcher intuition. Beyond a core panel of four frequently used CAF markers, this analysis identified seven additional markers that substantially improved the resolvability of clusters. Whether the resulting combination of 11 CAF-specific markers is sufficient to resolve CAF states, as observed with the 17 CAF-specific markers used here, remains to be verified experimentally.

Applying the FPP to PDAC tumors, we addressed open questions in CAF biology. We find that the LRRC15+/FAPα-high alternative myCAF metacluster (MC4) does not overlap with the αSMA-high/CD90-high contractile myCAF metacluster (MC2) (Fig. 2E), indicating LRRC15+ CAFs as a distinct myofibroblastic sub-state, not a subset of canonical αSMA+ myCAFs. Given that αSMA+ CAFs restrain PDAC growth while FAPα+ and LRRC15+ CAFs promote it,^6,21,25^ our data suggest that these are not simply different labels for the same activated state but rather mark divergent, functionally opposed myofibroblastic programs. This distinction has direct implications for CAF-targeted therapies, which may benefit from specifically targeting LRRC15+/FAPα+ populations.

The correlations between αSMA and CD90, pimonidazole and Ly6C, as well as pimonidazole and 2-NBDG signal across tumors (Sup. Fig. 5B) reproduce the previously reported relationship between two myofibroblast markers,^28^ enrichment of iCAFs in hypoxic areas,^15,31^ and the established link between hypoxia and glucose uptake,^35^ respectively. This validates the panel internally and supports confidence in less established correlations that we report here, such as LRRC15 and FAPα, or Ly6C and Ly6A. These correlations also exist at the mRNA level in autochthonous PDAC models (Sup. Fig. 5C).

Most markers show graded, not discrete, expression across PhenoGraph clusters (Fig. 2B-D). This argues against a strict trichotomous model of CAF identity; instead, iCAFs, myCAFs, and apCAFs may represent extremes along a continuum, with individual CAFs positioned variably between them.^11,12^ FPP’s resolution captures this continuum: it detects substructure even within larger, less polarized CAF populations, such as the Ly6A-high versus Ly6A-low clusters within Ly6C+ CAFs, a marker profile consistent with the Pi16+ universal fibroblast concept.^36^ This illustrates the value of high-dimensional flow cytometry profiling for capturing this continuum rather than forcing cells into a small number of discrete bins.

CAFs are highly plastic.^11^ *In vitro* experiments further demonstrate the FPP’s capacity to track plasticity outcomes, i.e., changes in fibroblast state induced by environmental stimuli. TGFβ stimulation polarized PSCs, collapsing basal PSC heterogeneity from five to two major clusters, accompanied by the loss of stem-like markers and the gain of myofibroblastic markers (Fig. 4F, G). While Cytokines+Hypoxia treatment enriched a cluster with more inflammatory characteristics and reduced clusters with myofibroblastic characteristics, it left overall PSC heterogeneity intact. This asymmetry could reflect differences in polarization strength between the two stimuli, or bias in the 2D culture model toward myofibroblastic states. It also shows that the FPP resolves stimulus-specific states even in the presence of the stiffness-driven myofibroblastic baseline imposed by 2D culture.

More complex responses were observed in the 3D culture model. Here, organoid co-culture and organoid CM upregulated inflammatory markers in PSCs, while still retaining some myofibroblastic characteristics (Fig. 3E). While metaclustering was sufficient to resolve the overall state changes induced by organoid-derived factors, differences between organoid-induced inflammatory- and myofibroblastic-leaning states were resolved at the fine cluster level. These data indicate that the mixed, multi-factor signal from organoid co-culture or CM can be separated into at least two independent inputs that can drive different arms of the CAF polarization program. These may be driven by IL1α and TGFβ, both of which are secreted by PDAC organoids.^14,28^ Notably, organoid-derived factors induce a CD90- state in PSCs (Fig. 3I), reminiscent of the recently described TGFβ-induced, EGFR-driven CAFs^28^. In addition, organoid co-culture but not CM induced a Ly6C/Ly6A-high state low in most other markers (Fig. 3C-E; Sup. Fig. 6B-D), similar to what we observed *in vivo* (Fig. 2B-D). These data indicate that direct contact with tumor cells induces stem-like characteristics in fibroblasts.

The FPP opens several directions. The emergence of spectral sorters allows sorting of FPP- defined populations, for example, the LRRC15+/FAPα-high state identified here, enabling direct functional testing of whether this population is functionally distinct from canonical αSMA+ myCAFs, thereby closing the phenotype-function gap left by transcriptomic profiling alone. In addition, it enables the differentiation of Ly6C/Ly6A-high stem-like/universal CAFs from other Ly6C/Ly6A-high CAFs by their low expression of all other markers, allowing further mechanistic and functional interrogation. Although we demonstrated the use of the FPP in the context of PDAC, fibroblasts play important roles in other cancer types as well as in other physiological and pathophysiological conditions across tissues, making our FPP well suited to study fibroblast heterogeneity in other solid tumors, wound healing, inflammation, and fibrosis. With 28 markers described here, there is still room for researchers to include other markers based on their question or model, with the major limitation being the availability of suitable fluorophore-conjugated antibodies. For example, this may include markers for recently described reticular CAFs identified in some PDAC tumors.^38^ Lastly, while our panel was designed with spectral flow cytometry in mind, it is also amenable to conventional 5-laser flow cytometers with a few modifications.

## Limitations of this study

Despite advances in resolving phenotypic CAF states, the FPP has limitations. As an antibody-based method, it resolves heterogeneity only along axes for which validated antibodies are available and included in the panel. It cannot detect unanticipated markers or states as unbiased transcriptomic profiling can. Therefore, the FPP complements scRNA-sequencing as a method for routine, affordable hypothesis testing; it does not replace it. Moreover, the scarcity of suitable commercially available antibodies for reported fibroblast state markers somewhat limits FPP’s capacity and resulted in several polyclonal antibodies being included in the panel, which may introduce lot-to-lot variability that could affect intra- and inter-lab reproducibility. In addition, sample fixation to assess intracellular markers precludes the use of FPP-stained cells for downstream studies. This may be overcome by including CAF state reporters, such as the IL6- EGFP and αSMA-DsRed dual reporter system we recently developed.^15^ The panel we report here was developed for readily available murine samples and cell lines, and a similar FPP will need to be developed for human samples and cell lines to support translational relevance. In addition, Ly6C and Ly6A have no human orthologs, precluding validation of the stem-like CAF population reported here in clinically relevant specimens. Finally, tissue dissociation required for flow cytometry analysis results in the loss of spatial information. The inclusion of pimonidazole as a hypoxia marker helps mitigate this with respect to tumor perfusion, but how phenotypically distinct CAF subsets identified with the FPP relate to other cell types and to each other will require building a similar panel for multiplex immunostaining and imaging.

### Key Resources

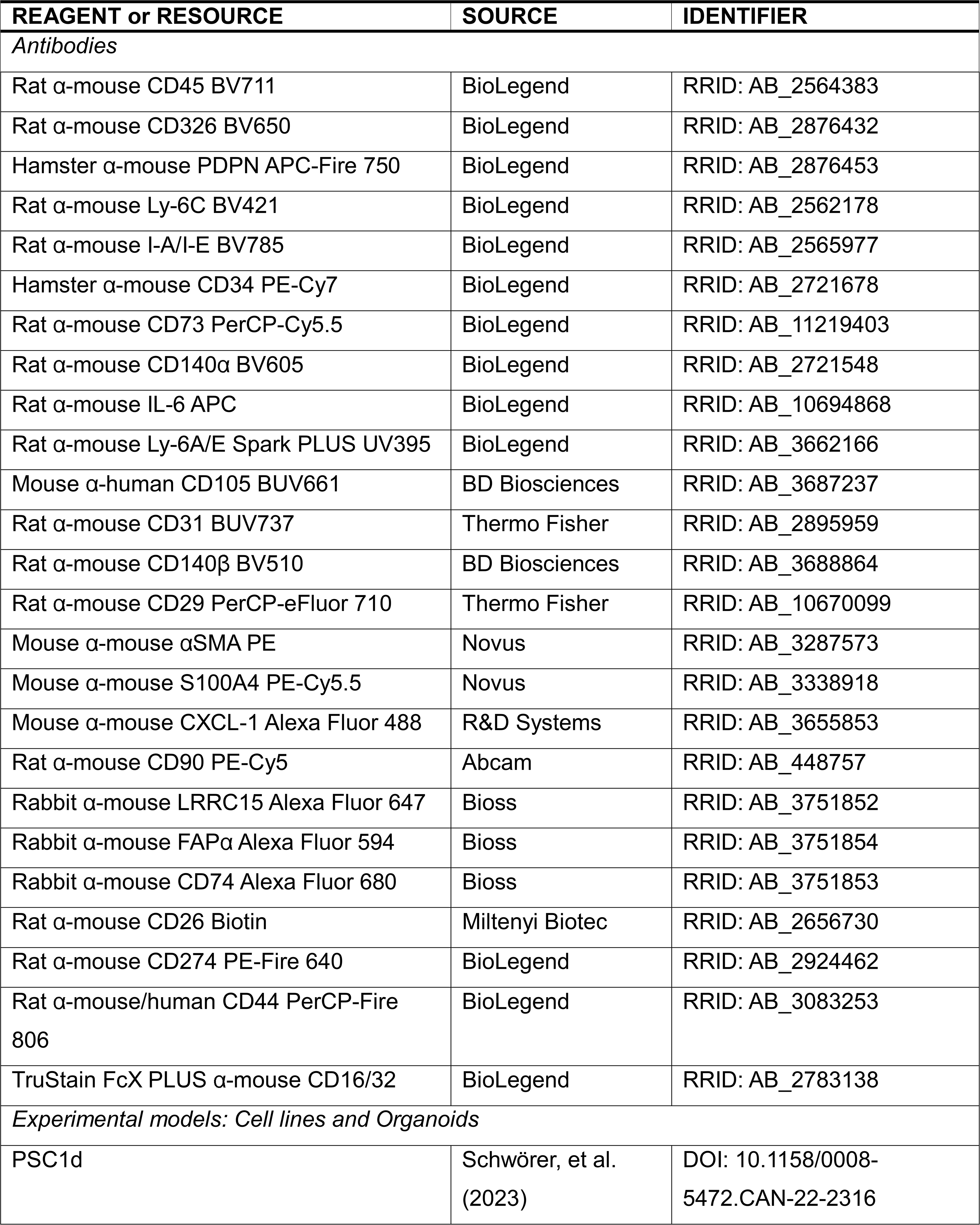
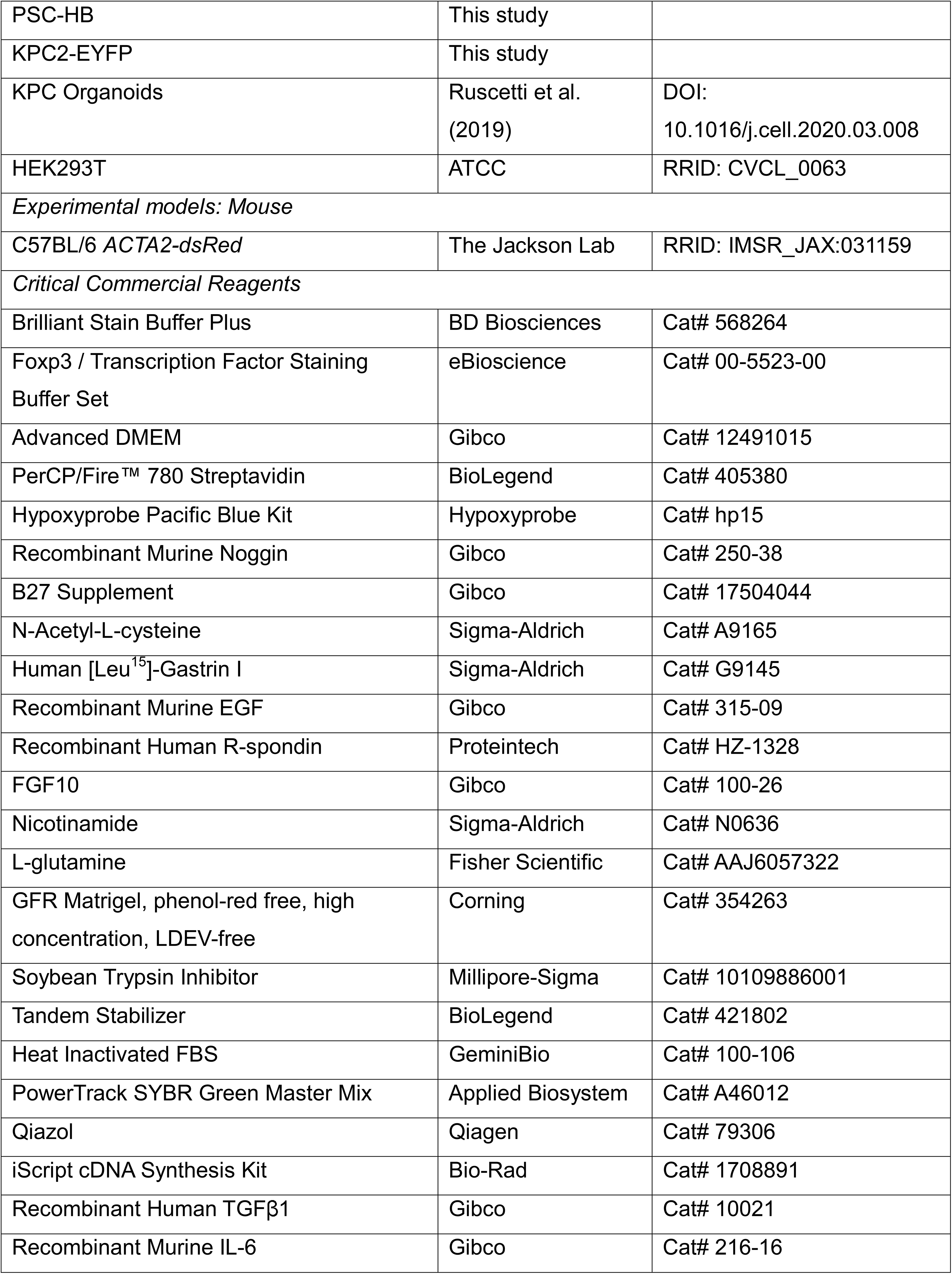
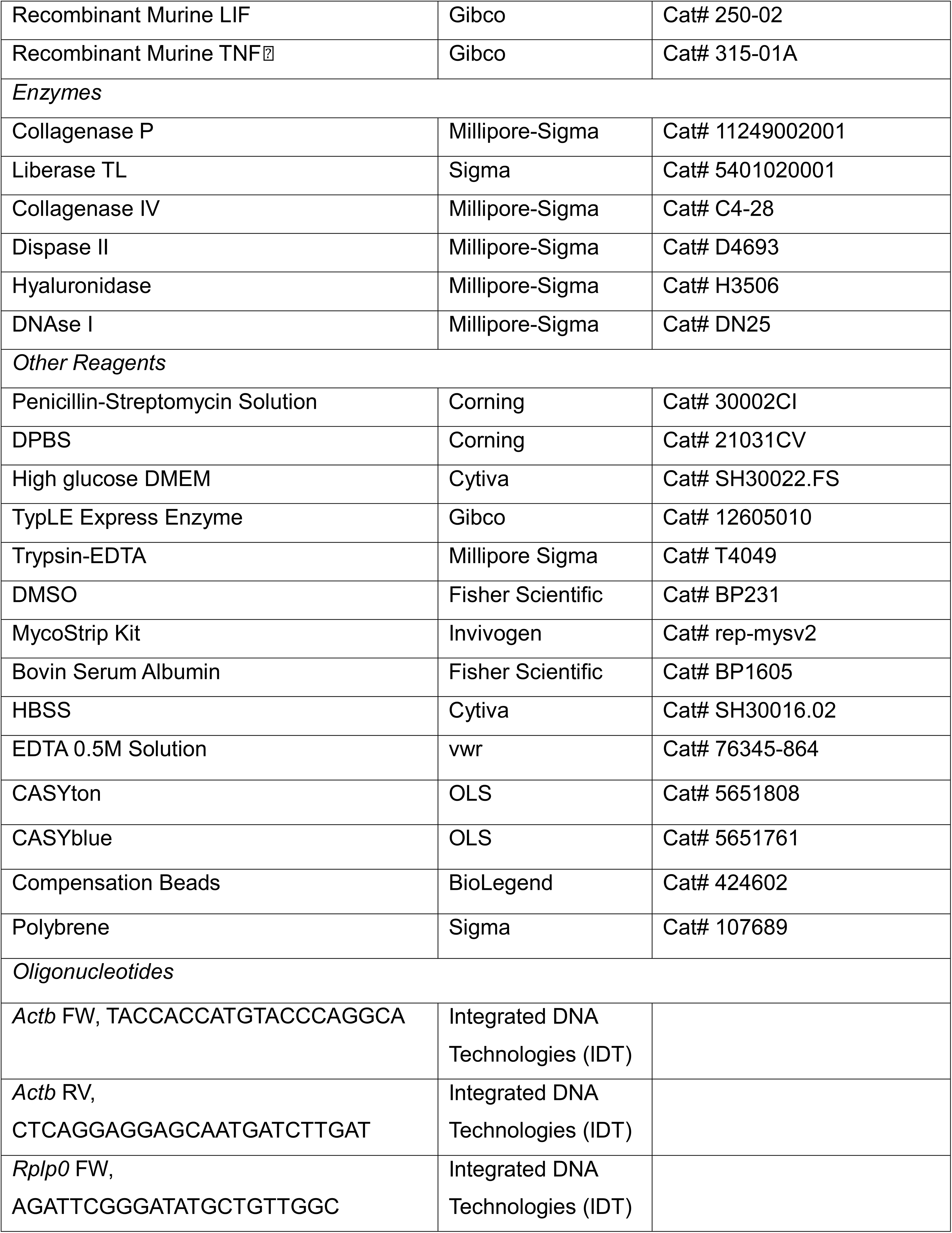
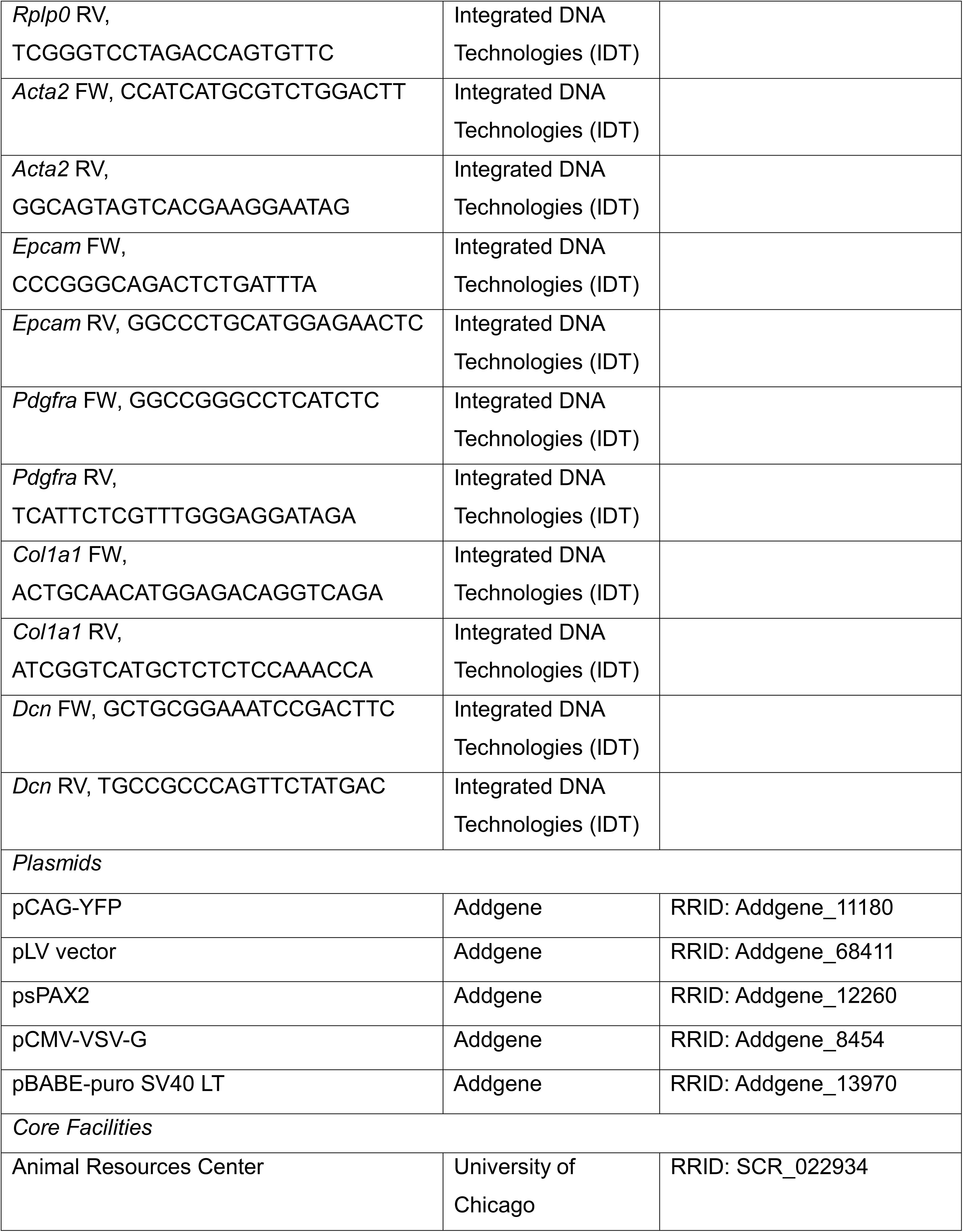
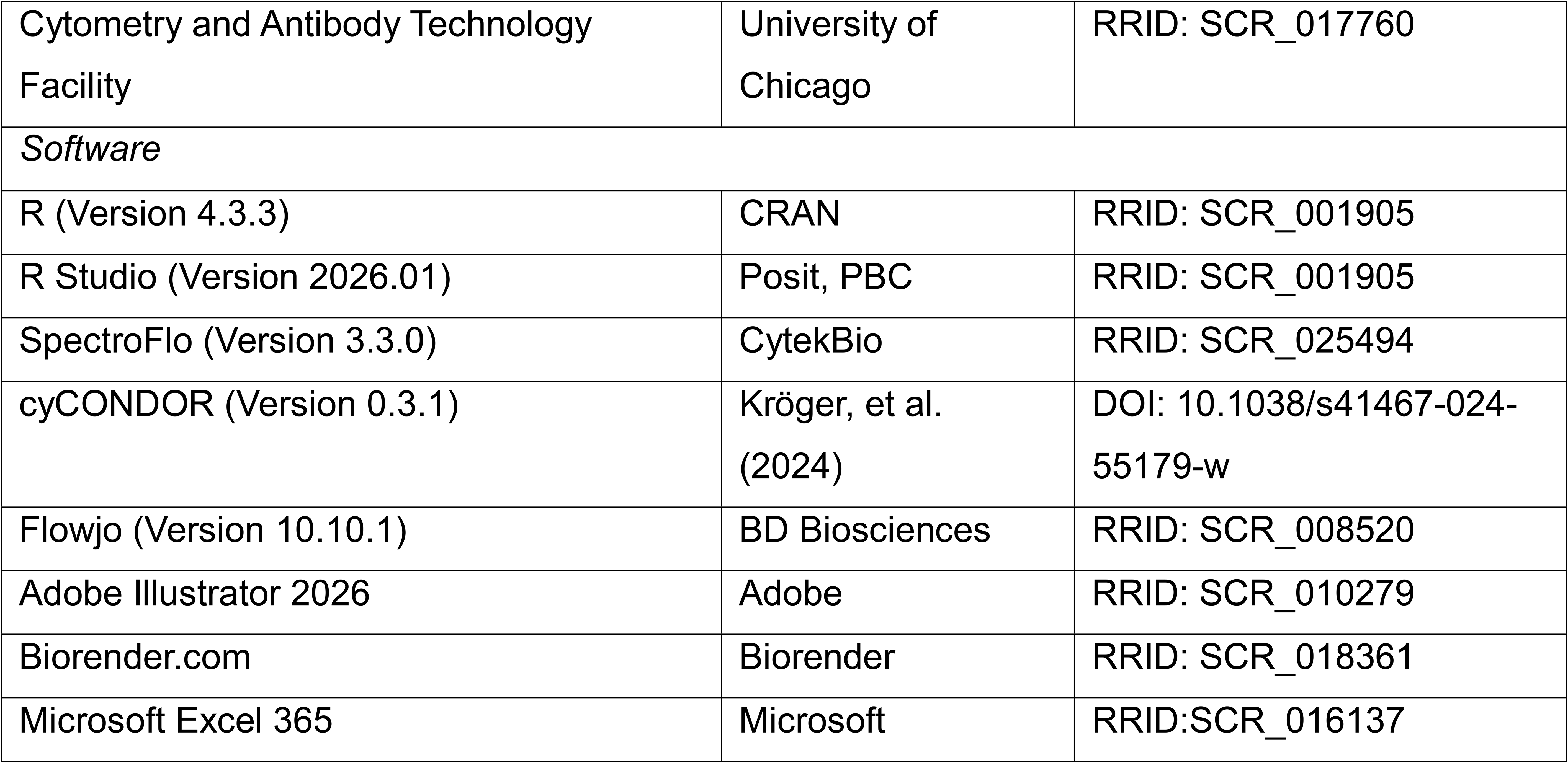

## Methods

### Experimental Models

#### Animal Models

Acta2-DsRed C57BL/6 mice were purchased from Jackson Labs and housed at the University of Chicago Animal Resources Center vivarium on a 12-hour day/night cycle with chow and water *ad libitum*. Mice wildtype for the Acta2-DsRed transgene were utilized for experiments at approximately 10–12 weeks of age. All experimental animal procedures were conducted in accordance with National Institutes of Health guidelines and were approved by the University of Chicago’s Institutional Animal Care and Use Committee under protocol ACUP #72708.

#### in vitro Models

KPC2 (*Kras*^LSL-G12D/+^;*Trp53*^LSL-R172H/+^;*Pdx1-Cre*) cells and organoids were a gift from the laboratory of Scott Lowe at Memorial Sloan Kettering Cancer Center (New York, NY) and previously described.^39^ KPC59 (*Kras*^LSL-G12D/+^;*Trp53*^fl/fl^;*Pdx1*-Cre) cells were a gift from the laboratory of Alexander Muir at The University of Chicago (Chicago, IL) and previously described.^40^ KPC2-EYFP cells were generated as part of this study as described below. Both KPC2 and KPC2-EYFP cells were cultured on plastic tissue culture plates coated with 0.1% rat tail collagen I in PBS. PSC1d cells were previously isolated and characterized,^15^ and PSC-HB cells were isolated as previously described,^15^ immortalized with SV40-LT, and characterized in this study. All cell lines were maintained under BSL-2+ conditions in high-glucose DMEM supplemented with 10% FBS and 1% penicillin/streptomycin at 37 °C in a water-saturated atmosphere of 5% CO_2_. KPC organoids were cultured in suspension in 100 µL of growth factor-reduced Matrigel (GFR-Matrigel) at 8.3 mg/mL in advanced DMEM/F12 supplemented with the following: 1% penicillin/streptomycin, 2 mM glutamine, 1X B27 supplement, 50 ng/ml EGF, 100 ng/ml Noggin, 100 ng/ml hFGF10, 10 nM Leu^15^-Gastrin I, 1.25 mM N-acetylcysteine, 10 mM nicotinamide, and 0.5 µg/mL R-spondin. Cell lines and PDAC organoids were routinely tested for mycoplasma using the MycoStrip kit (Invivogen) and were used for <15 passages and <6 passages, respectively. Cell lines and PDAC organoids were cryopreserved in liquid nitrogen in their respective growth media supplemented with 10% DMSO.

### Generation of KPC2-EYFP Cell Line

KPC2-EYFP cells were generated by lentiviral transduction of parental KPC2 cells with pLV- EF1 :EYFP. This plasmid was generated by Gibson assembly of the EYFP cassette from pCAG-YFP into the pLV vector. Lentiviral particles were produced in 293T cells using psPAX2 and pCMV VSV-G packaging plasmids. Media was changed the next day, and viral supernatant was collected 48 hours later, passed through a 0.45 μm nylon filter, and used for transduction in the presence of 8 μg/ml polybrene. Successfully transduced cells were selected by sorting EYFP+ cells.

### Generation of *in vivo* PDAC Model

To generate PDAC tumors, 200,000 KPC2-EYFP cells or 2 confluent wells of KPC organoids grown in 24-well plates (∼50,000 cells) were suspended in 20 µL Cultrex 3D Culture Matrix at 4.15 mg/mL and implanted into the tail of pancreata of wild-type *Acta2*-DsRed mice. KPC2- EYFP tumors were allowed to grow up to 5 weeks, and KPC organoid tumors were allowed to grow up to 4 months. Mice were monitored daily to assess their health status, and tumor growth was monitored via palpation and ultrasound.

### PDAC organoid and PSC co-culture

PDAC organoids were isolated by mechanically and enzymatically digesting their surrounding Matrigel droplet using TrypLE, and each well of organoids was split at a ratio of 1:6. For organoid:PSC co-culture, 1 well equivalent of organoids and 80,000 PSCs were resuspended in 50 µL of GFR-Matrigel each, then mixed together and transferred into a 24-well TC plate. For the 2D, 3D, and conditioned media (CM) groups, 80,000 PSCs were cultured directly in a 24- well TC plate (2D) or resuspended in 100 µL of GFR-Matrigel (3D and CM). Conditioned media were generated by allowing PDAC organoids to reach confluence, after which their growth media were replaced with high-glucose DMEM supplemented with 5% FBS, and the organoids were cultured for an additional 3 days. The conditioned DMEM was recovered, centrifuged at 800 x g for 5 minutes, and filtered using a 0.2 µm syringe filter. In each assay, 16 wells per group were seeded simultaneously and fed 500 µL of DMEM supplemented with 5% FBS, or, for the CM group, 500 µL of 50% PDAC organoid-conditioned media diluted with 50% fresh DMEM supplemented with 5% FBS. After 3 days of culturing, the PSCs were lifted using TrypLE (2D) or recovered from the Matrigel by mechanical dissociation and enzymatic digestion. To digest the Matrigel, enzymes were added to each well to a final concentration of 0.2 mg/mL collagenase IV, 0.2 mg/mL DNase I, and 0.5 mg/mL Dispase II, and the Matrigel dome was mechanically disrupted using a 1000 µL tip. The mixture was incubated for 25 minutes in a TC incubator and then again mechanically disrupted using a 1000 µL tip. Three wells worth of cells were combined into a 15 mL conical tube with 10 mL ice-cold HBSS, and the mixture was incubated on ice for 10 minutes. After 10 minutes, the samples were centrifuged at 600 × g for 5 minutes at 4 °C. The resulting cell pellets were resuspended in HBSS with 2 mM EDTA (HBSS+EDTA) and stained for flow cytometry analysis as described below.

### 2D culture and treatment

PSCs were seeded in 12-well TC plates at 120,000 cells per well with 1 mL of supplemented DMEM and allowed to recover overnight in a TC incubator. The following day, corresponding treatments were added to achieve the following final concentrations: PBS + 0.1% BSA (vehicle), 10 ng/mL TGFβ, or a cytokine cocktail at 2 ng/mL each of LIF, TNFα, and IL1α. Plates treated with the cytokine cocktail were moved to a hypoxia chamber at 37 °C in a water-saturated atmosphere with 1% O_2_. After 3 days of culture under their respective treatment conditions, the cells were lifted with TrypLE and stained for flow cytometry analysis as described below.

### qPCR

RNA was isolated from KPC59 and PSC-HB cells with Qiazol, and cDNA was generated from 1 ug RNA with the iScript cDNA Synthesis Kit. Quantitative real-time PCR (qPCR) analysis was performed in technical triplicate using 1:20-diluted cDNAs and 0.1 µM forward and reverse primers, together with PowerTrack SYBR Green, on a QuantStudio 7 Flex (Applied Biosystems). Gene expression was quantified in Microsoft Excel 365 as relative expression ratio using primer efficiencies calculated by a relative standard curve. The geometric mean of the endogenous control genes *Actb* and *Rplp0* was used as the reference sample. Primer sequences are listed in the Key Resources table.

### Isolation of PDAC-derived CAFs

Tumor-bearing mice were euthanized via isoflurane overdose followed by cervical dislocation. Tumor tissues were isolated and minced using surgical scissors. Minced tissue was then placed in a 50 mL conical tube with 6 mL of an enzymatic digestion solution containing 0.8 mg/mL Dispase II, 0.5 mg/mL Collagenase P, 0.1 mg/mL Liberase TL, 0.1 mg/mL DNAse I, 0.4 mg/mL Hyaluronidase, and 0.02 mg/mL Trypsin inhibitor, 2.5 mM CaCl_2_, diluted in HBSS. Samples were then transferred to C-tubes and processed using program 37C_m_TDK1_2 on a gentleMACS Octo dissociator with heaters (Miltenyi Biotec). Enzymatic digestion was quenched by adding 14 mL of cold HBSS, and the samples were centrifuged at 500 × g for 5 minutes at 4°C. The resulting cell pellets were resuspended in 5 mL ACK lysis buffer and incubated at RT for 5 minutes. Lysis was quenched by adding 10 mL of HBSS+EDTA, and the samples were centrifuged at 500 × g for 5 minutes at 4°C. The resulting cell pellets were resuspended in 5 mL HBSS+EDTA and filtered through a 70 µm mesh filter. The mesh filter was washed with 5 mL HBSS+EDTA, and the cell concentrations were determined using a CASY cell analyzer. Samples were stained for flow cytometry analysis as described below.

For studies using metabolic probes, 1 hour before tumor tissue isolation, mice were injected i.p. with pimonidazole HCl (Pimo), diluted in sterile saline, at 60 mg/kg body weight. Once a single-cell suspension was obtained, cells were incubated with 40 µM 2-NBDG in PBS at a concentration of 1 million cells per 100 µL for 15 minutes at 37°C in the dark. 2-NBDG uptake was quenched by washing twice with 10X volumes of ice-cold HBSS+EDTA.

### Antibody Staining and Flow Cytometry

#### Antibody and Viability Dye Titration

Optimal antibody and viability dye concentrations were determined by titrating each antibody at 0.125X, 0.25X, 0.5X, 1X, and 2X, where X is the manufacturer’s recommended concentration. PDAC tumors were mechanically and enzymatically dissociated as described above to generate a single-cell suspension. Cells were counted and adjusted to 2 million cells/100µL in HBSS+EDTA. Cells were Fc-blocked with anti-mouse TruStain FcX™ PLUS at 1:100 for 15 minutes at RT. For extracellular markers, cells were stained with a single antibody at a specified concentration, diluted in HEBs (HBSS with 2 mM EDTA and 5% BSA), supplemented with 1X Brilliant Stain Buffer Plus (BSB), for 1 hour at 4 °C in the dark. Cells were then fixed and permeabilized using the FoxP3 Transcription Factor Staining Buffer Set, supplemented with 1X Tandem Stabilizer (TS) for 45 minutes, and stored at 4 °C in the dark until data collection. For intracellular markers, after FC blocking, cells were fixed and permeabilized as described above. After washing with 1X permeabilization buffer, cells were resuspended in the intracellular antibody diluted in 1X permeabilization buffer and incubated overnight at 4°C in the dark. The following day, the cells were washed twice with 2X volumes of 1X permeabilization buffer, resuspended in HEBs, and stored at 4°C in the dark until data collection. For the viability dye, 2 million cells/concentration were heat-killed by incubation at 70 °C for 5 minutes. Half of these were stained with the viability dye for 15 minutes at RT in the dark, and the other half remained unstained. Staining was quenched with HEBs, and the cells were fixed and permeabilized as described above. Data were acquired on a 5L Cytek Aurora spectral flow cytometer at the University of Chicago Cytometry and Antibody Technology Core. Staining indices for each antibody were determined using the Stain Index plugin for FlowJo. For the final concentrations of each antibody, see Supplemental Table 1.

#### Sample Staining for Flow Cytometry Analysis

Cells were counted and adjusted to a concentration of 2 million cells/100 µL in HBSS+EDTA. Samples were incubated in HBSS+EDTA with Zombie NIR [1:2,000] and FC block [1:100] at a concentration of 2 million cells/100 µL for 15 minutes at RT in the dark. Viability staining was quenched by adding 2X volumes of HEBs and washing the samples for 5 minutes at 600 × g at 4 °C. An extracellular antibody cocktail was prepared by diluting the appropriate antibodies in HEBs supplemented with BSB. Samples were brought to a concentration of 2 million cells/100 µL of extracellular antibody cocktail and incubated for 1 hour at 4 °C in the dark. After staining, samples were washed twice in 2X volumes of HEBs to remove excess antibody. For intracellular staining, stained cells were fixed and permeabilized as described above. An intracellular antibody cocktail was prepared in 1X permeabilization buffer supplemented with TS. Samples were washed in 1X volume of permeabilization buffer to remove fixative, resuspended in intracellular antibody cocktail at 2 million cells/100 µL, and incubated overnight (>12 hours) at 4 °C in the dark. The following day, the samples were washed twice with 1 mL permeabilization buffer, resuspended in HEBs supplemented with TS, and stored in the dark at 4°C until data collection. In parallel to cell staining, spectral control samples were prepared. Cell-based controls were prepared as described above to unmix Zombie NIR. To extract sample autofluorescence, 100,000 cells from each sample were aliquoted, fixed, and permeabilized. For antibody staining, compensation beads were singly stained with the fluorophore-conjugated antibodies. For spectral unmixing of EYFP, KPC2 and KPC2-EYFP cells were fixed and permeabilized as described above. Data collection was conducted at The University of Chicago Cytometry and Antibody Technology Core lab on a 5L Aurora spectral flow cytometer.

#### Flow Cytometry Data Analysis

All data processing was conducted in R using cited, publicly available packages and using OpenBLAS integration for parallel multithreaded matrix calculations on a Windows PC, unless otherwise specified.

Unmixed FCS files were exported from SpectroFlo, and in Flowjo the samples were preprocessed by first gating on singlets, removing outlier events with FlowAI using the standard exclusion values.^37^ CAFs were gated on as depicted in Figure 1C, and then randomly downsampled to the lowest individual sample cell count for equal representation. Preprocessed CAF events were exported as .FCS files for analysis in R. Using the R package *premessa*, .FCS files were parsed, and housekeeping parameters, specifically FSC, SSC, SSC-B, Time, autofluorescence, and the viability dye (Zombie NIR), were removed to exclude them from all downstream analyses. *cyCONDOR* was used for general analysis.^41^ Dimensionality reduction (UMAP) and clustering were performed with *uwot* and *Rphenograph*. Protein marker expression projections were plotted using outlier clamping for values below the 1st percentile and above the 99th percentile and point sorting to ensure a maximized dynamic range and prevent visual occlusion of high/low expressing events, respectively.

To evaluate intra-sample homogeneity of variances in cell clustering as a function of the number of markers, Bartlett’s test of sphericity was applied to the individual single-cell correlation matrix for the selected markers in R. Based on the field’s established gating strategy, a core set of four mandatory markers (Ly6C, MHC II, PDPN, and CD140α) was fixed. A combinatorial search was conducted over all possible combinations for the remaining markers to identify the combination of markers that maximized Bartlett’s statistic, which was selected as the minimal set of markers. Spearman’s rank correlation coefficients were computed for all unique marker pairs. A Fisher Z-transformation was applied to normalize the distributions across samples. One-sample *t*-tests were conducted on the Z-scores for each pair against a null hypothesis of zero correlation. To control the false discovery rate (FDR) from multiple testing across all pairs, *p*-values were adjusted using the Benjamini-Hochberg procedure, with statistical significance defined at an FDR-adjusted *q*<0.05.

#### scRNA-sequencing Correlation analysis

Single-cell RNA-sequencing (scRNA-seq) data from KPC mice were previously collected, filtered, and normalized by Elyada et al.^27^ and are available at Gene Expression Omnibus under the accession number GSE129455. Downstream analysis was performed using R 4.5.1 and the following R packages: “readr_2.2.0’, “scCorr_0.1”, “Seurat_5.5.0”, “org.Mm.eg.db_3.21.0”, “biomaRt_2.64.0”, “ggpubr_0.6.3”, “ggplot2_4.0.3”, “RColorBrewer_1.1-3”, “dplyr_1.2.1”, and “AnnotationDbi_1.70.0”. *CreateSeuratObject()* was used to prepare the gene expression matrix for the standard Seurat workflow, wherein highly variable features were identified using *FindVariableFeatures()*, counts were scaled using *ScaleData()*, and principal component analysis (PCA) was performed using *RunPCA()*. Cell clusters were calculated and identified using *FindNeighbors()* and *FindClusters()* with parameters determined by *ElbowPlot()*. *RunUMAP()* was used to generate UMAPs, and *DimPlot()* was used to visualize the result. CAFs were identified as described by Elyada et al. and isolated for downstream analysis using *subset()*. *AddModuleScore()* was used to calculate the "HALLMARK_HYPOXIA" (MSigDB 2026.1) signature score.

To reduce error and noise caused by the RNA-seq “dropout” effect, scCorr, a k-partitioning method that merges transcriptomically similar cells, was employed as described.^42^ The optimal k-value was iteratively determined to maximize the number of clusters with ≥10 cells. Pearson correlation coefficients with 95% confidence intervals were calculated between two genes or between a gene and a signature using *cor.test()* within the merged scCorr cells. Correlations were visualized using *ggplot()*.

## Acknowledgements

The authors thank Dr. Kim Cardenas (BioLegend) for consultation on panel design.

## Author Contributions

Conceptualization, K.M.F. and S.S.; Data Curation, K.M.F. and M.S.D.; Formal Analysis, K.M.F., M.S.D., S.S.; Funding Acquisition, K.M.F. and S.S.; Investigation, K.M.F., M.S.D., S.E.I., D.M., M.N. and S.L.; Methodology, K.M.F., M.S.D., A.S. and S.S.; Project Administration, K.M.F. and S.S.; Resources, K.M.F. and S.S.; Software, K.M.F. and M.S.D.; Supervision K.M.F. and S.S.; Validation, K.M.F., M.S.D., S.E.I., and D.M.; Visualization, K.M.F., M.S.D., and S.S.; Writing – Original Draft, K.M.F., S.S.; Writing – Review and Editing, K.M.F., M.S.D., and S.S.

## Supplemental Table 1. Antibody dilutions

List of antibodies and their dilution factor used in this study.

---

**Supplemental Figure 1.**
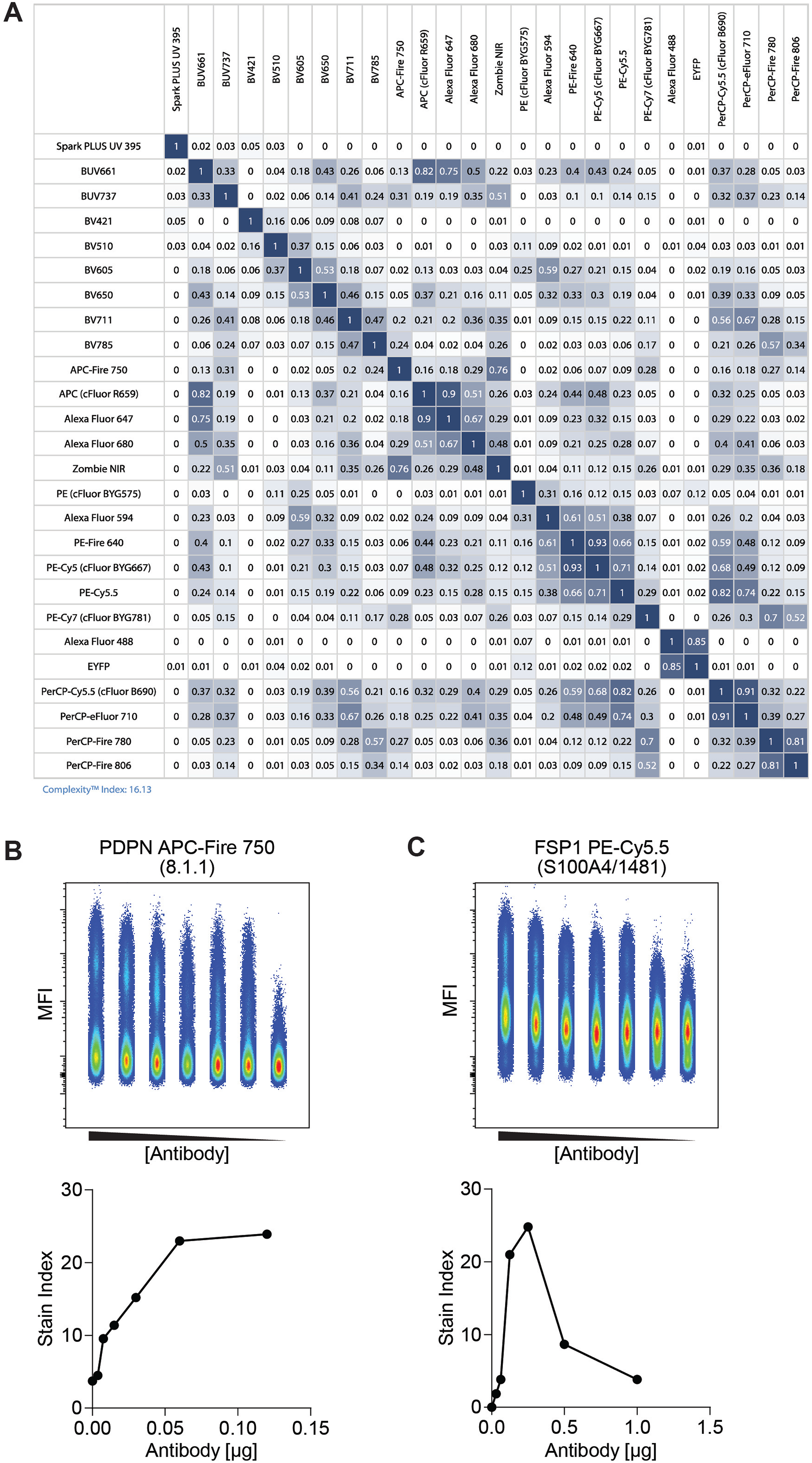
Design and optimization of core fibroblast state profiling panel. **(A)** Similarity matrix of spectral signatures for core fibroblast state profiling panel. **(B, C)** Representative plots of 7-point serial antibody titrations (top) and stain index calculated for each antibody concentration (bottom) for PDPN (B) and FSP1 (C).

**Supplemental Figure 2.**
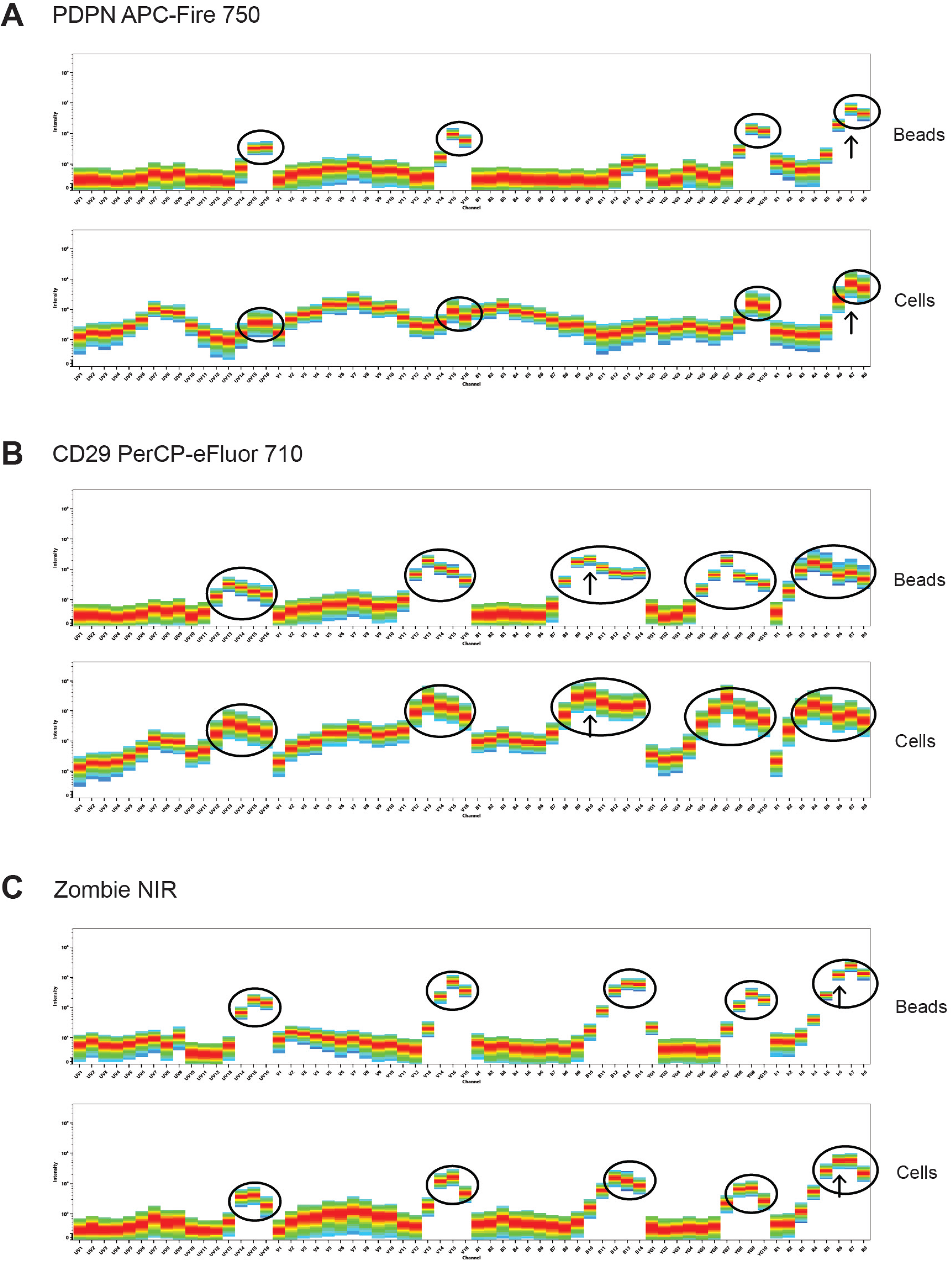
Comparison of cell- and bead-based spectral signatures. For optimal spectral unmixing, the spectral signature of each fluorophore-antibody pair was compared between single-stained polystyrene compensation beads (top) and cells derived from tumor tissues (bottom). Representations of spectrographs of protein-based tandem fluorophores (A) and (B) and of the amine-binding dye Zombie NIR (C). Circles highlight major signature peaks, and arrows indicate the principal peak.

**Supplemental Figure 3.**
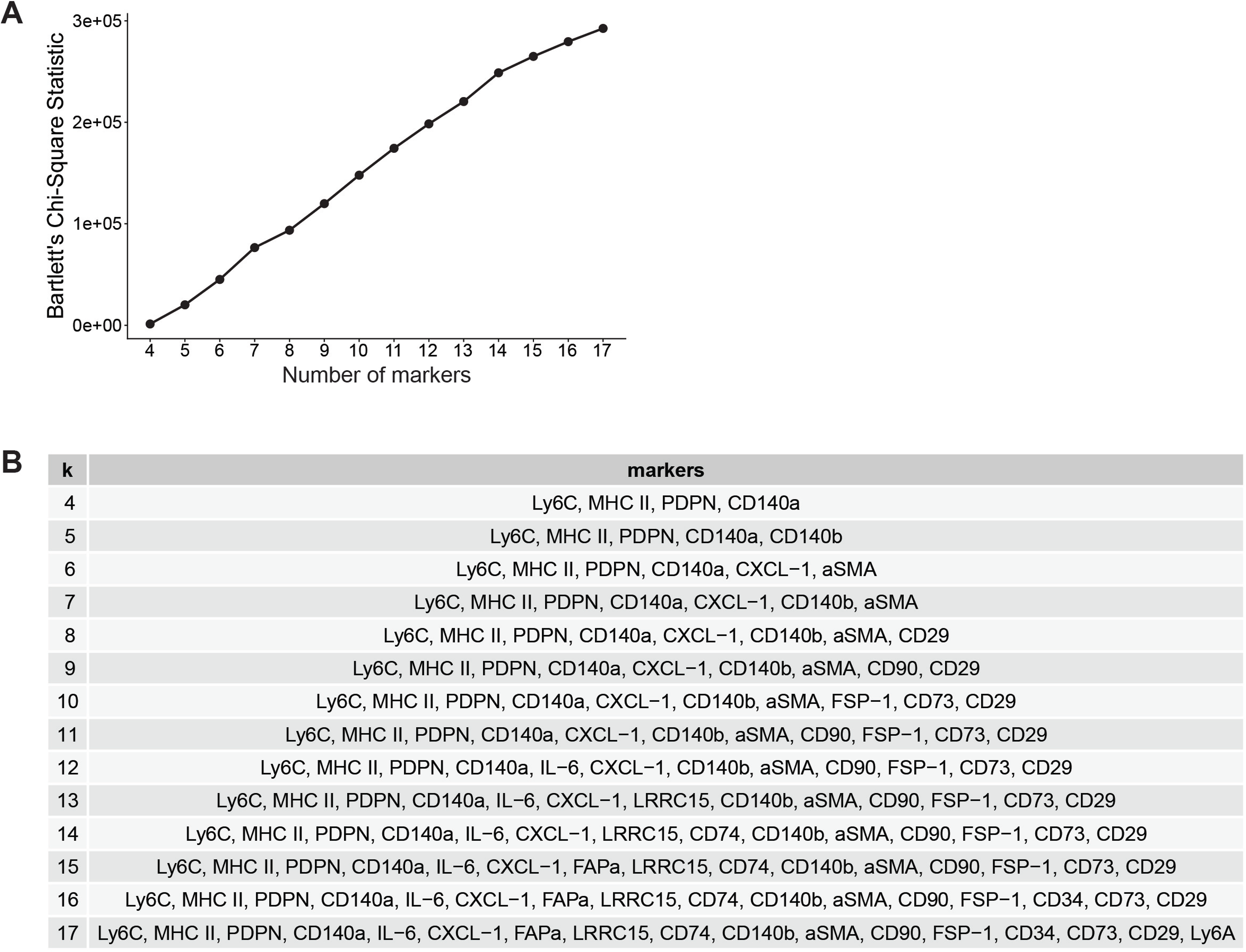
Identification of inter-sample variance. Starting from a population of cells that were negative/low for Zombie NIR, negative for CD45, CD326, CD31, and EYFP, and positive for PDPN and/or PDGFRα, we iteratively calculated the homogeneity of variances of cell clusters as a function of the number of markers utilizing Bartlett’s test. **(A)** Plot of the number of markers included as a function of Bartlett’s chi-square statistic. **(B)** Table of markers that provided the most stable variance for each iteration of k number of markers

**Supplemental Figure 4.**
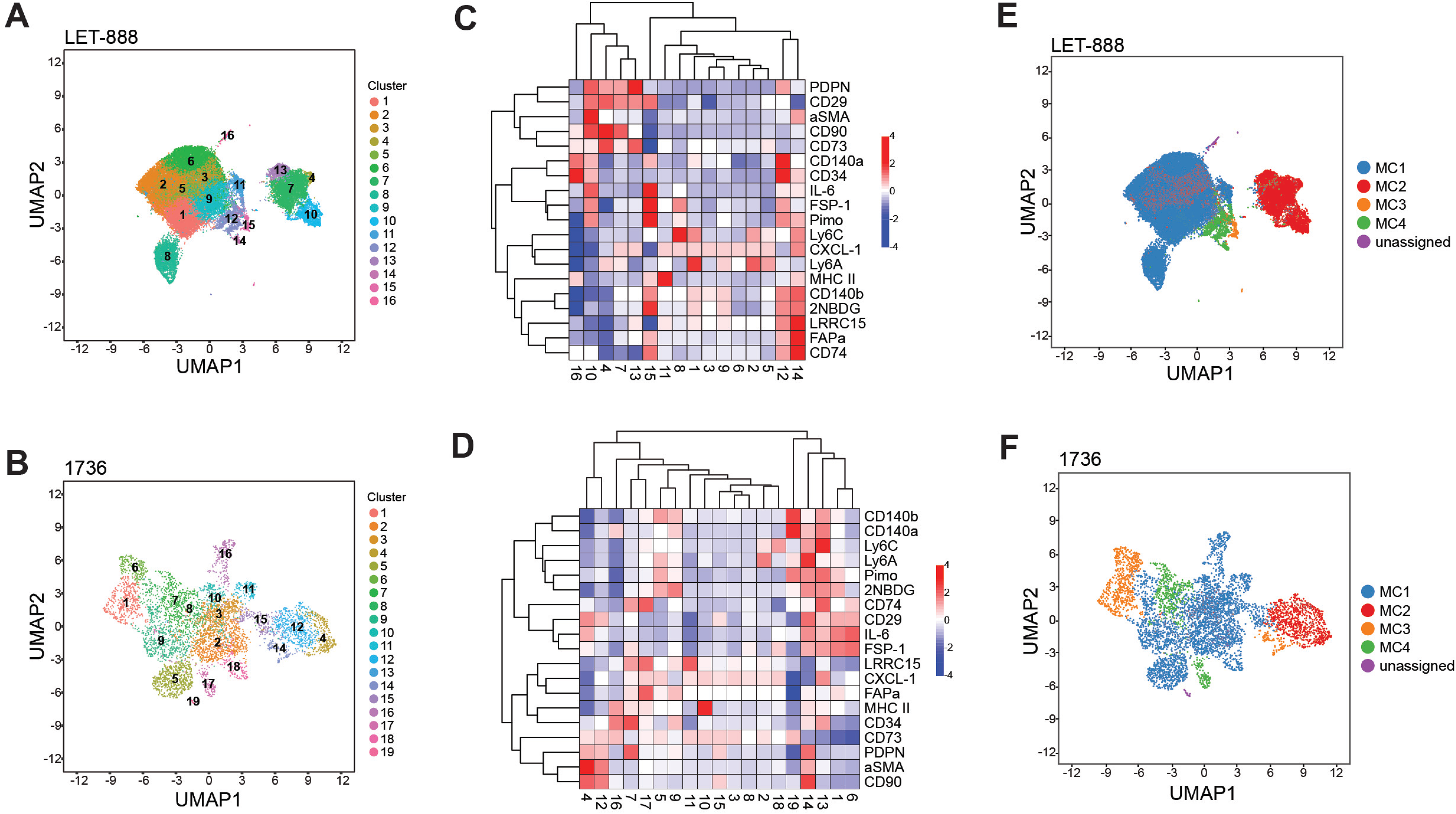
Fibroblast state profiling panel demonstrates robust inter-sample consistency in PhenoGraph clustering and cluster similarity in PDAC tumors. **(A, B)** Paired UMAPs with PhenoGraph clustering for two independent biological replicates. Tumors LET-888 and 1736 are shown. **(C, D)** Paired heatmaps of marker expression across PhenoGraph clusters with hierarchical clustering for two independent biological replicates. Tumors LET-888 and 1736 are shown. **(E, F)** Paired UMAPs with metaclustering for two independent biological replicates. Tumors LET- 888 and 1736 are shown.

**Supplemental Figure 5.**
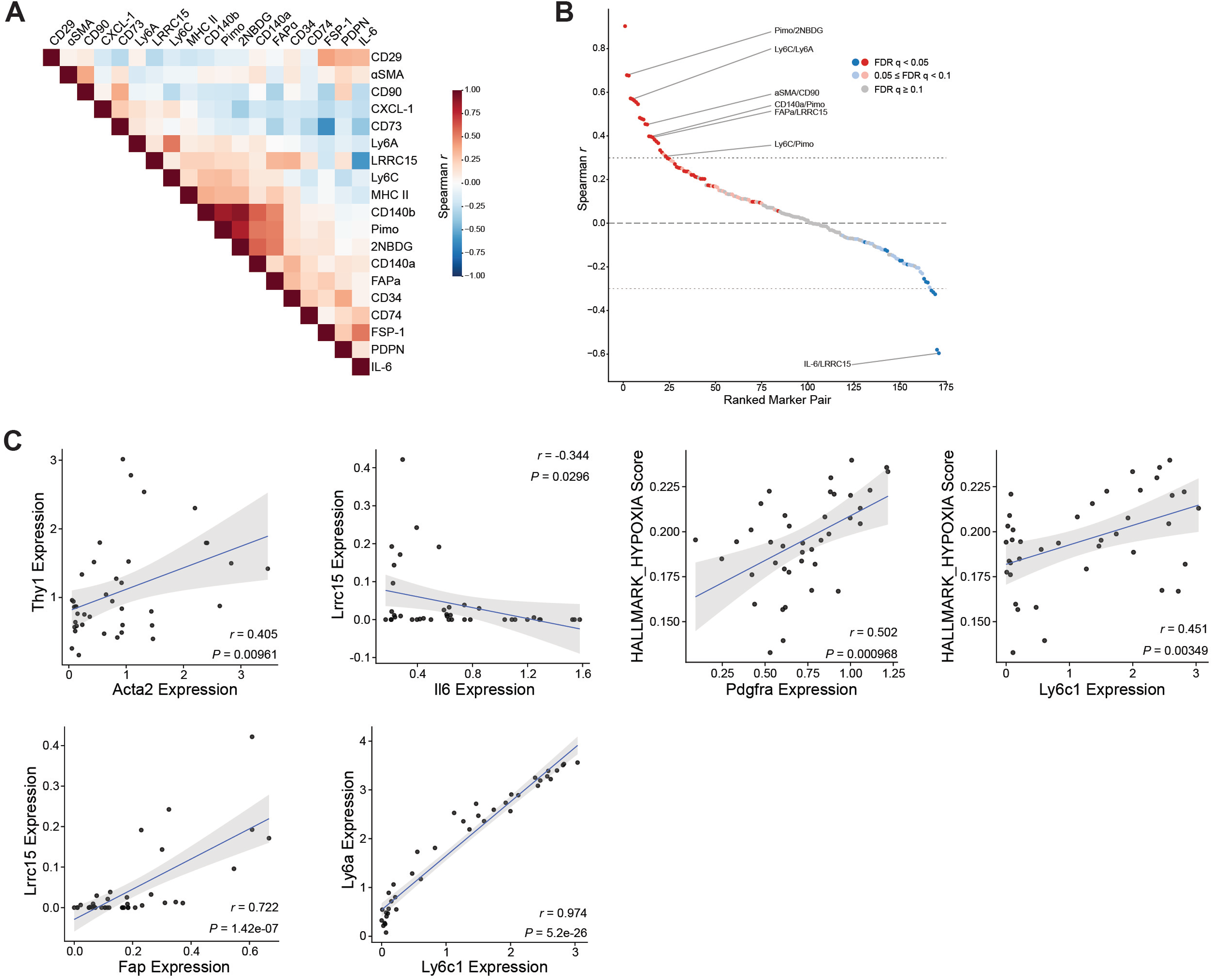
Fibroblast state profiling panel exhibits inter-marker correlation signatures consistent with transcriptomic analyses. **(A)** Heatmap of Spearman *r* values. **(B)** Plot of ranked marker pairs as a function of Spearman *r* values for all possible marker combinations across replicate tumors. **(C)** Correlation plots of scRNA-seq between select significant marker pairings identified using the FPP.

**Supplemental Figure 6.**
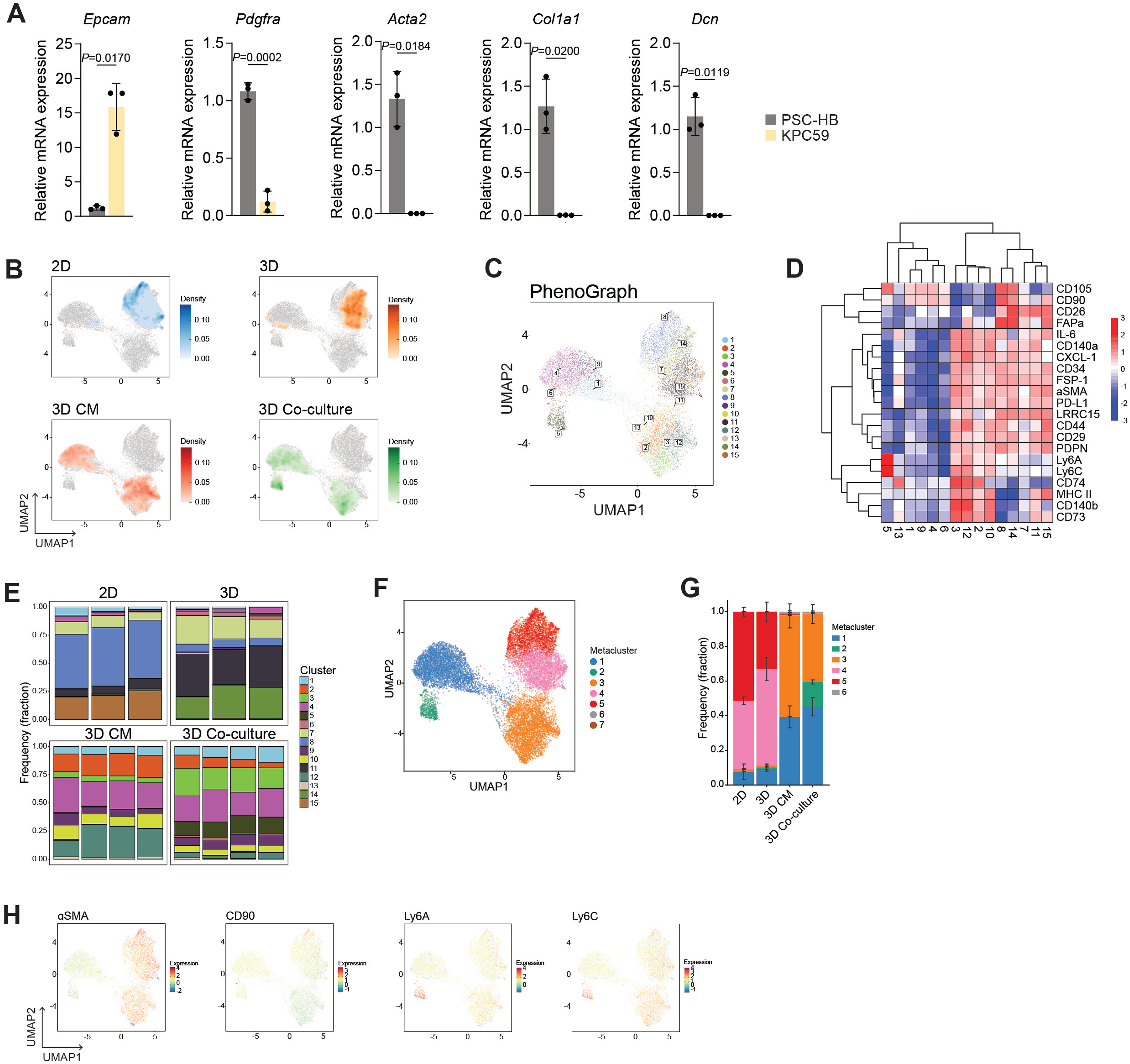
PDAC organoid co-culture exhibits differential direct and indirect effects on PSC-HB fibroblast cell state. **(A)** Bar charts of relative gene expression for the indicated genes in PSC-HB and KPC59 cells to validate PSC-HB’s fibroblast nature. *n*=3 biological replicates. Data represent mean +/- SD. *P*-values were calculated by Student’s *t*-test. **(C)** UMAPs colored by cell density, faceted by culture condition. **(C)** UMAP with PhenoGraph clustering projected. **(D)** Heatmap of marker expression across PhenoGraph clusters with hierarchical clustering. **(E)** Bar charts of composition of each PhenoGraph cluster as a function of sample grouping. **(F)** UMAPs of select CAF markers with marker expression overlay. **(G)** UMAPs overlaid with the expression of select CAF markers. **(H)** Bar charts of composition of each metacluster as a function of sample grouping. *n*=3 (2D/3D), *n*=4 (3D CM, 3D Co-culture) biological replicates. Data show mean +/- SD.

